# Virus-associated Inflammation Imprints an Inflammatory Profile on Long-lived Monocyte-derived Macrophages in the Human Liver

**DOI:** 10.1101/2024.01.31.578178

**Authors:** Juan Diego Sanchez Vasquez, Shirin Nkongolo, Daniel Traum, Samuel C. Kim, Deeqa Mahamed, Aman Mehrotra, Anjali Patel, Diana Chen, Scott Fung, Anuj Gaggar, Jordan J. Feld, Kyong-Mi Chang, Jeffrey J. Wallin, Harry L.A. Janssen, Adam J. Gehring

## Abstract

Chronic liver injury triggers the activation and recruitment of immune cells, causing antigen-independent tissue damage and liver disease progression. Tissue inflammation can reshape macrophage composition through monocyte replacement. Replacement of tissue macrophages with monocytes differentiating in an inflammatory environment can potentially imprint a phenotype that switches the liver from an immunotolerant organ to one predisposed to tissue damage. We longitudinally sampled the liver of chronic hepatitis B (CHB) patients with active liver inflammation starting antiviral therapy. Antiviral therapy suppressed viral replication and liver inflammation, which coincided with decreased myeloid activation markers. Single-cell RNA sequencing mapped peripheral inflammatory markers to a monocyte-derived macrophage population, distinct from Kupffer cells, with an inflammatory transcriptional profile. The inflammatory macrophages (iMacs) differentiated from blood monocytes and established a long-lived population. The iMacs retained their core transcriptional signature, consistent with trained immunity, resulting in a population of macrophages primed for inflammation potentially driving progressive liver disease.

## Introduction

Macrophages (MΦs) play a pivotal role in tissue immunity, representing one of the first lines of defense against pathogens. They regulate tissue damage via pathogen clearance and secretion of cytokines, chemokines, and growth factors to avoid unchecked inflammation(*1*). The MΦ pool is highly heterogeneous, consisting mainly of self-renewing embryonic-derived MΦs that seed the organs during fetal development, and monocyte-derived MΦs recruited from the blood upon tissue damage or infection(*2, 3*). Mouse studies have revealed monocyte recruitment and imprinting is tissue-specific, allowing them to differentiate into either long-lived or short-lived MΦs(*4–6*).

During acute infections, reshaping the macrophage population through an influx of monocyte-derived macrophages improves pathogen clearance and return to baseline (*7*). However, over 800 million people live with chronic liver disease (CLD), which is characterized by persistent inflammation, leading to over 2 million deaths per year from cirrhosis and hepatocellular carcinoma (HCC) (*8, 9*). In mouse models of liver inflammation, MΦs contribute to disease progression by producing inflammatory cytokines that activate surrounding intrahepatic cells, driving inflammation and tissue injury, causing progressive fibrosis that impairs liver regeneration(*8*). These inflammatory events can be ameliorated by MΦ depletion in experimental models, underscoring their central role in tissue regulation (*3, 10*).

MΦ-mediated inflammation presents a stark contrast to the steady state environment of the liver, which is one of tolerance and immune-suppression. The tolerogenic environment is maintained via IL-10 production from Kupffer cells (KCs), embryonically-derived MΦs of the liver (*11, 12*). This suggests that during chronic liver disease, the KC niche is being reshaped by monocyte-derived MΦs that are more inflammatory. This could change the dynamics of immune regulation in the liver and contribute to chronic liver damage, as has been observed in mouse models (*10, 13–15*)

Understanding how inflammation reshapes liver MΦ composition in humans has been restricted by the ability to collect longitudinal tissue samples within a time frame relevant to inflammation dynamics. However, patients with chronic Hepatitis B virus (HBV) infection that present to the clinic with active liver damage are started on antiviral therapy. Antiviral therapy suppresses HBV replication and stops liver damage within 6 months. Once started, antiviral treatment can be life-long to maintain suppression of viral replication. Withdrawal of therapy leads to HBV reactivation and potentially life threatening liver inflammation that is characterized by a MΦ signature(*16–20*). Using longitudinal liver fine-needle aspirates (FNAs), we captured dynamic changes in cellular composition and activation during the first 6 months of therapy (*21*). By combining liver FNA sampling in patients starting antiviral therapy with single-cell RNA sequencing (scRNAseq), we achieved a resolution necessary to define heterogeneous MΦ populations and their activation status in the liver of patients with chronic hepatitis B (CHB). This led to the identification of an inflammatory, long-lived monocyte-derived MΦ population unique to inflamed livers. These monocyte-derived MΦs were imprinted with an inflammatory profile that was distinguishable even after 4 years of antiviral therapy.

## Results

### Serum markers of myeloid activation map to liver MΦs

To investigate cellular dynamics of liver inflammation and damage in CLD, we recruited 11 CHB patients with active ongoing liver damage as evidence by elevated levels of serum alanine aminotransferase (ALT) represented as fold increase over normal values (* upper limit of normal (*ULN >1)). Patients started antiviral treatment with tenofovir alafenamide (TAF) (25mg daily), which reduced liver damage and HBV replication(*22*) (Fig. 1A). We collected liver FNAs at study entry and 12 and 24 weeks post-TAF initiation and performed scRNAseq in five patients (red lines) to link inflammatory biomarkers in the serum to activation of immune cells in the liver (Fig 1B). This allowed us to compare intrahepatic immune profiles during liver injury (baseline) and as inflammation resolved (weeks 12 and 24).

**Figure 1.**
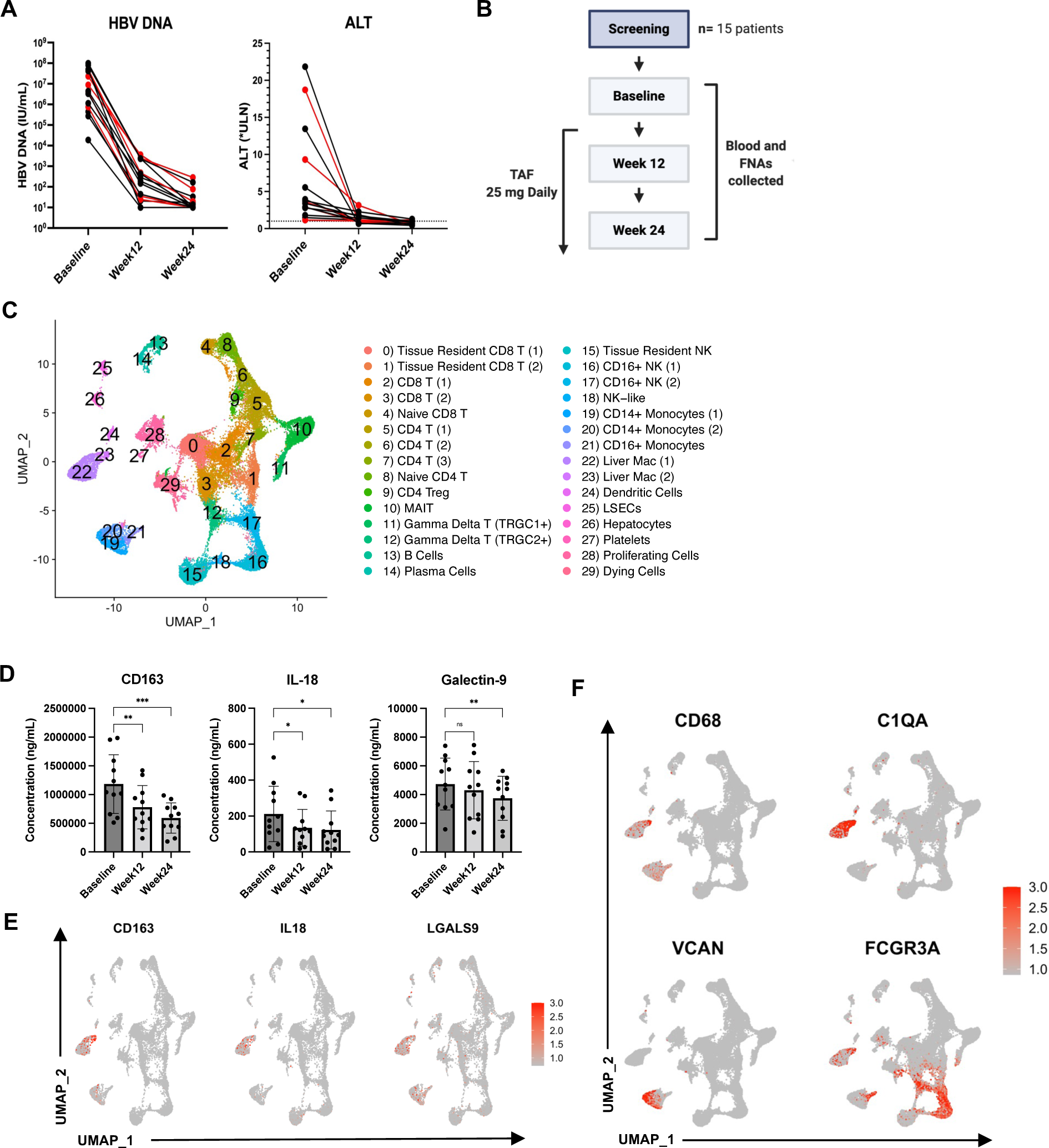
Study Design and identification of hepatic populations at single-cell resolution from liver FNAs. **A)** HBV DNA and alanine aminotransferase (ALT) (displayed as fold-change of upper limit of normal) in blood over time, patients whose samples were sequenced are highlighted in red (**n=**5). **B)** 11 chronic hepatitis B (CHB) patients with elevated ALT started nucleoside analogue therapy with tenofovir alafenamide (TAF) 25mg/d. At baseline and after 12 and 24 weeks of therapy, blood and liver fine-needle aspirates (FNAs) were collected. Longitudinal FNAs from 5 patients were subjected to single-cell RNA sequencing (scRNAseq). **C)** Clustering and annotation of 41,829 cells from human livers for 5 patients across 3 timepoints (baseline, week12 and week24). UMAP dimensionality reduction identified 30 clusters. **D)** Luminex data from plasma immune profile of all patients across time. **E)** Feature plots depicting single-cell gene expression of individual genes detected by the Luminex assay. **F)** Feature plots depicting single-cell gene expression of liver myeloid cells. *P*values determined by repeated-measures one-way ANOVA (\**P* <0.05, \*\**P* <0.005, \*\*\**P* <0.001).

We identified 30 distinct cell clusters from 41,829 cells across the three longitudinal time points, which showed a similar distribution among the different donors (Fig. 1C and Suppl. Fig. 1A and 1B). We previously demonstrated that the myeloid activation marker soluble CD163 (sCD163) was significantly increased during liver damage across multiple stages of HBV infection(*23*). Analysis of serum immune markers in our patient cohort confirmed elevated levels of sCD163 in the serum of patients with hepatitis that significantly declined after 12 weeks of TAF therapy. Consistent with the decline in sCD163, additional myeloid activation markers IL-18, and galectin-9 also significantly decreased after starting therapy (Fig 1D). Using the transcriptome data, we mapped these markers to the monocyte and MΦ clusters, defined by the expression of *CD68* (Monocyte and MΦ marker), *C1QA/B/C* (MΦ marker), *VCAN* (classical monocyte marker) and *FCGR3A* (non-classical monocyte marker) (Fig. 1E and 1F)(*24–26*). These data validated our previous observations(*23*) and implicate monocytes and MΦ as potential key players in the pathogenesis of liver injury in CHB.

### Inflammatory MΦs are phenotypically distinct from Kupffer cells

Because serum markers suggested myeloid involvement, CD68+ cells were subclustered for further analysis. This led to the identification of 2 classical monocyte clusters (expression of *VCAN, LYZ*, and *CD14*), one non-classical monocyte cluster (CD16+ monocytes; *FCGR3A* (CD16a)*, SIGLEC10* and *PECAM1*); and two MΦ clusters (Liver Mac (1) and Liver Mac (2); *C1QA* (complement component), *SLC40A1* (Ferroportin) and *MARCO*)), (Fig. 2A-2B)(*4, 5, 24–27*).

**Figure 2.**
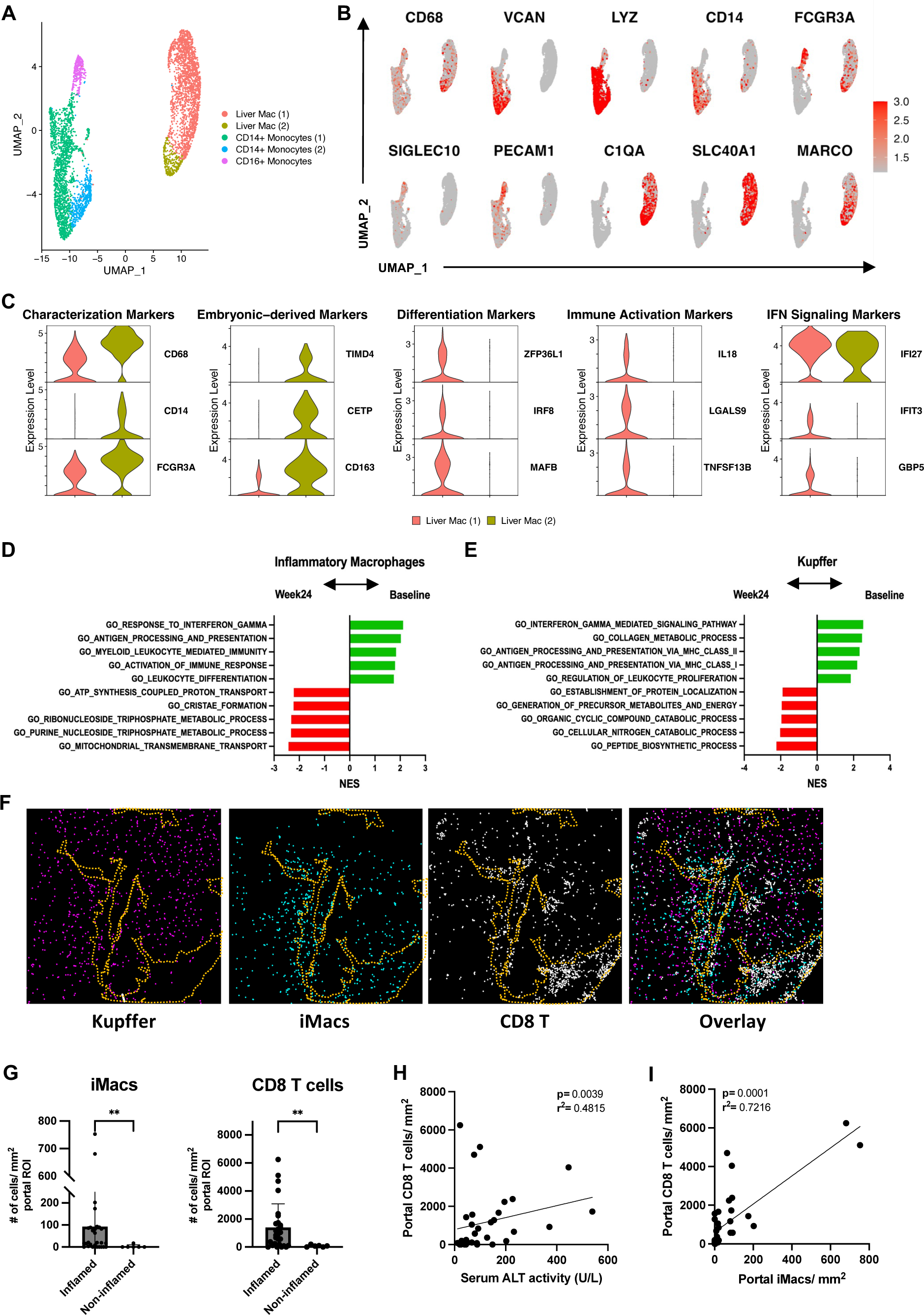
Identification and characterization of myeloid cells during liver inflammation. **A)** CD68+ clusters were re-clustered using UMAP dimensionality reduction for 5 patients across 3 timepoints (baseline, week12 and week24). **B)** Feature plots depicting single-cell gene expression of individual myeloid genes (scale: log-transformed gene expression) **C)** Violin plots of Mac-defining genes. All selected genes have an adjusted p value < 0.05. **D, E)** Pathway analysis (GSEA) of differentially expressed genes in inflammatory Macs **(D)** and Kupffer cells **(E)** at baseline versus week24 of TAF treatment. Pathway enrichment is expressed as the Normalized Enrichment Score (NES) for multiple comparisons. **F)** IMC depicting liver Macs and CD8 T cell co-localization in inflamed liver of CHB patients. Portal region outlined in yellow dotted lines. **G)** Quantification of liver Macs and CD8 T cells/mm^2^ in portal area ROI between inflamed (**n=**28) and non-inflamed (**n=**6) liver tissue sections. *P* values determined by non-parametric Mann Whitney U test, (\**P* <0.05, \*\**P* <0.005, \*\*\**P* <0.001). **H and I)** Simple correlation analysis between portal CD8 T cells/ mm^2^ and **H)** iMacs/mm^2^ and **I)** serum ALT levels in CHB patients. Macs, macrophages; iMacs, inflammatory macrophages; IMC, imaging mass cytometry; ROI, region of interest.

Both MΦ clusters expressed *CD68* and *FCGR3A* (CD16a), but Liver Mac (1) could be distinguished by the lack of *CD14* expression (Fig 2C, first column). Liver Mac (2) expressed markers of embryonic derived MΦs such as *TIMD4*(*28, 29*)*, CETP*, and *CD163* (Fig 2C, second column). In contrast, Liver Mac (1) was enriched for markers of recent monocyte-to-MΦ differentiation (*ZFP36L1, IRF8,* and *MAFB*)(*4, 5, 30, 31*) (Fig 2C, third column), immune activation (*IL18, LGASL9* (Galectin-9) and *TNFSF13B (*BAFF))(*32–34*) (Fig 2C, fourth column), and interferon signaling (*IFI27, IFIT3 and GBP5*)(*35, 36*) (Fig 2C, fifth column). These data demonstrate the co-existence of multiple MΦ populations during active liver damage in CHB patients. Based on the transcriptional profiles, Liver MΦ (1) and Liver MΦ (2) will be referred to as inflammatory MΦ (iMacs) and Kupffer cells (KCs), respectively.

Consistent with transcriptional differences of each MΦ population, their predicted functional profiles differed by Gene Set Enrichment Analysis (GSEA)(*37*). GSEA was performed between baseline and week 24. Processes related to inflammation, interferon signaling, and differentiation were enriched in the iMacs at baseline (Fig. 2D). KCs were also enriched in IFN signaling and antigen presentation but differed by pathways associated with collagen metabolism and proliferation (Fig. 2E). These data underscore the heterogeneous nature of MΦs and how they may have unqiue responses during liver inflammation, despite residing in the same organ.

To validate differences in the transcriptional phenotype at the protein level, we performed imaging mass cytometry (IMC) on biopsies from CHB patients with (n=28) and without (n=6) liver inflammation. KCs, defined as CD68^+^CD16^+^CD14^+^, were evenly distributed throughout the tissue (Fig 2F). In contrast, iMacs characterized as CD68^+^CD16^+^CD14^-^ significantly clustered near the portal regions and co-localized with CD8 T cells in inflamed tissues compared to non-inflamed tissues (Fig. 2F, G and Suppl. Fig. 2A, B). The number of periportal CD8 T cells significantly correlated with ALT level (Fig 2H) as well as the iMac frequency in the portal area (Fig 2I). Localization to the portal area suggests iMacs were recently recruited to the liver. Co-localization with CD8 T cells, known to cause antigen-independent liver damage, implicates iMac the progression of CLD(*21*).

### The iMacs are unique to the inflamed liver

The IMC data suggested that iMacs are restricted to patients with active liver inflammation. To confirm this observation at the transcriptional level, we compared MΦs taken from the baseline sample in our study and compared them to MΦs from uninfected healthy(*25*) and cirrhotic(*13*) human liver scRNAseq datasets. We identified 5 distinct MΦ populations (Fig. 3A). All liver MΦs were characterized by the expression of *C1QA, SPI1* and *FCGR3A* (Fig. 3B). Since all the other scRNAseq data sets were collected from tissue digestion of liver biopsies, they recovered a higher number of MΦ that contributed to better defined clusters compared to liver FNAs. For this reason, the cluster previously defined as KCs revealed 3 distinct sub-MΦ clusters with distinct phenotypes, TIMD4^+^ MΦ, TIMD4^-^MARCO^+^ MΦ and TIMD4^-^MARCO^-^ MΦ, which were shared among the 3 types of livers (Fig 3C). Scar Associated MΦs (SAMacs) were largely restricted to the cirrhotic liver whereas iMacs were almost unique to the inflamed liver (Fig. 3A, 3C) and displayed a distinct transcriptional signature of inflammation and differentiation compared to all other MΦ populations (Fig 3D). These data demonstrate that liver inflammation gives rise to a population of MΦ not found in healthy individuals or cirrhotic livers.

**Figure 3.**
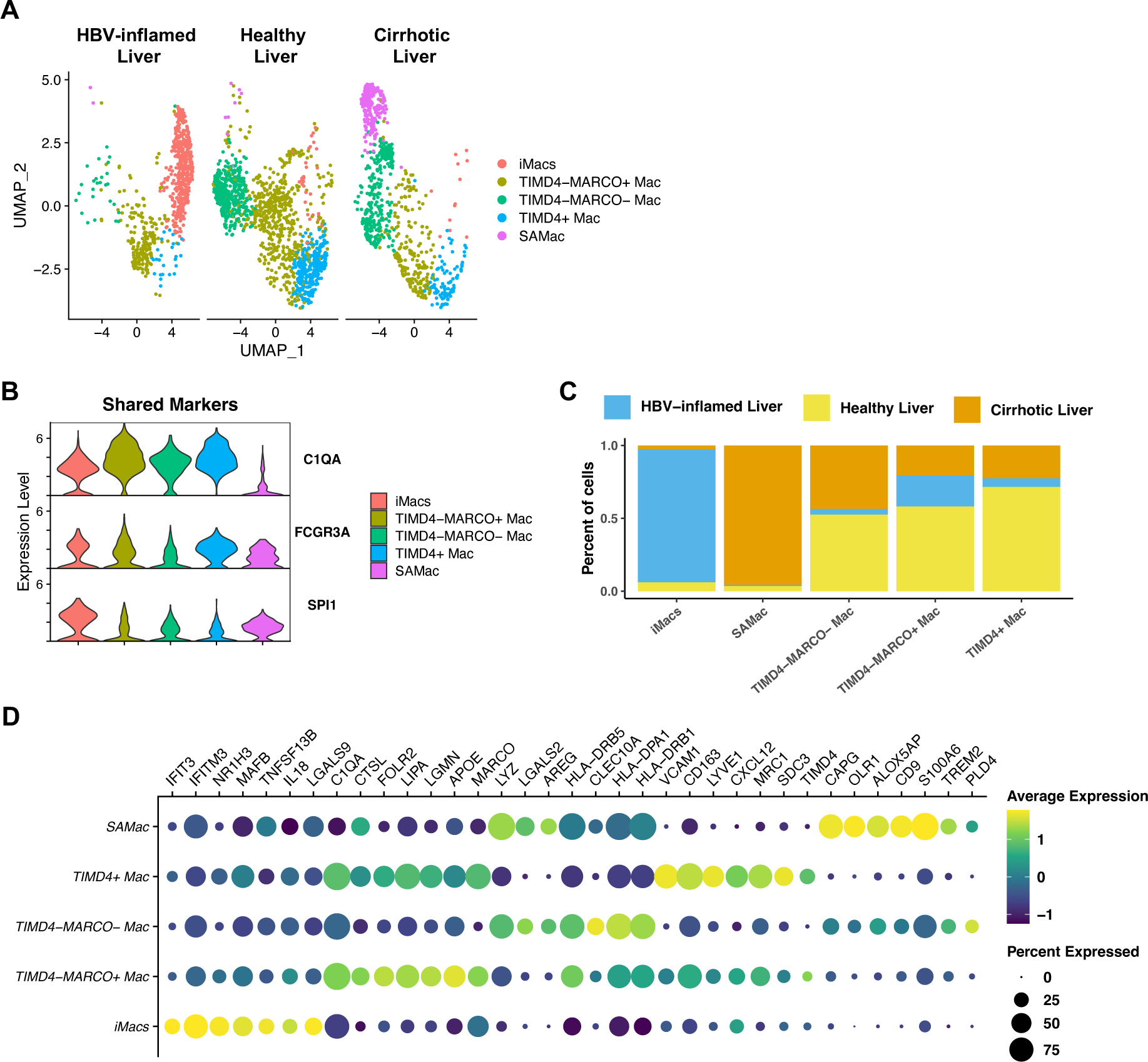
Comparison of inflammatory macrophages during HBV reactivation to healthy and cirrhotic livers at the single cell level. **A)** Clustering of Macs from healthy (**n=**5), cirrhotic (**n=**5) and HBV-inflamed (**n=**5) human livers using UMAP dimensionality reduction by liver condition. Cells from both healthy and cirrhotic livers were obtained from liver tissue digestion and sequenced with 10x Genomics 3’ v2, which explains why tissue digestion collected more Macs compared to the FNA approach. **B)** Violin plots of Mac-shared genes across 3 groups. All selected genes have an adjusted p value < 0.05. **C)** Proportions of liver Macs across 3 groups for each cluster. **D)** Representation of cluster defining genes, all selected genes have an adjusted p value <0.05. iMacs, inflammatory macrophages; Mac, macrophages; SAMacs, Scar-associated macrophages; HBV, hepatitis B virus.

### Trajectory analysis reveals monocyte-to-MΦ differentiation during liver inflammation

In animal models of liver disease, the KC pool becomes progressively replaced by infiltrating monocyte-derived MΦs, which can be more inflammatory(*10*). The transcriptional profile of iMacs suggests a similar pathway in the inflamed human liver (Fig 2C and 2D). Therefore, we assessed whether CD14+ monocytes displayed gene expression patterns consistent with differentiation into iMacs. Both CD14+ monocyte populations expressed *VCAN, CD14, FCN1* and *S100A8*, but distinguished from one another by the expression of *CCR2, CX3CR1, IFNGR1* and *IFNAR1* (Fig. 4A and 4B). GSEA analysis of cluster CD14+ monocytes (1) indicated up-regulation of pathways associated with myeloid differentiation during liver inflammation (Fig. 4C).

**Figure 4.**
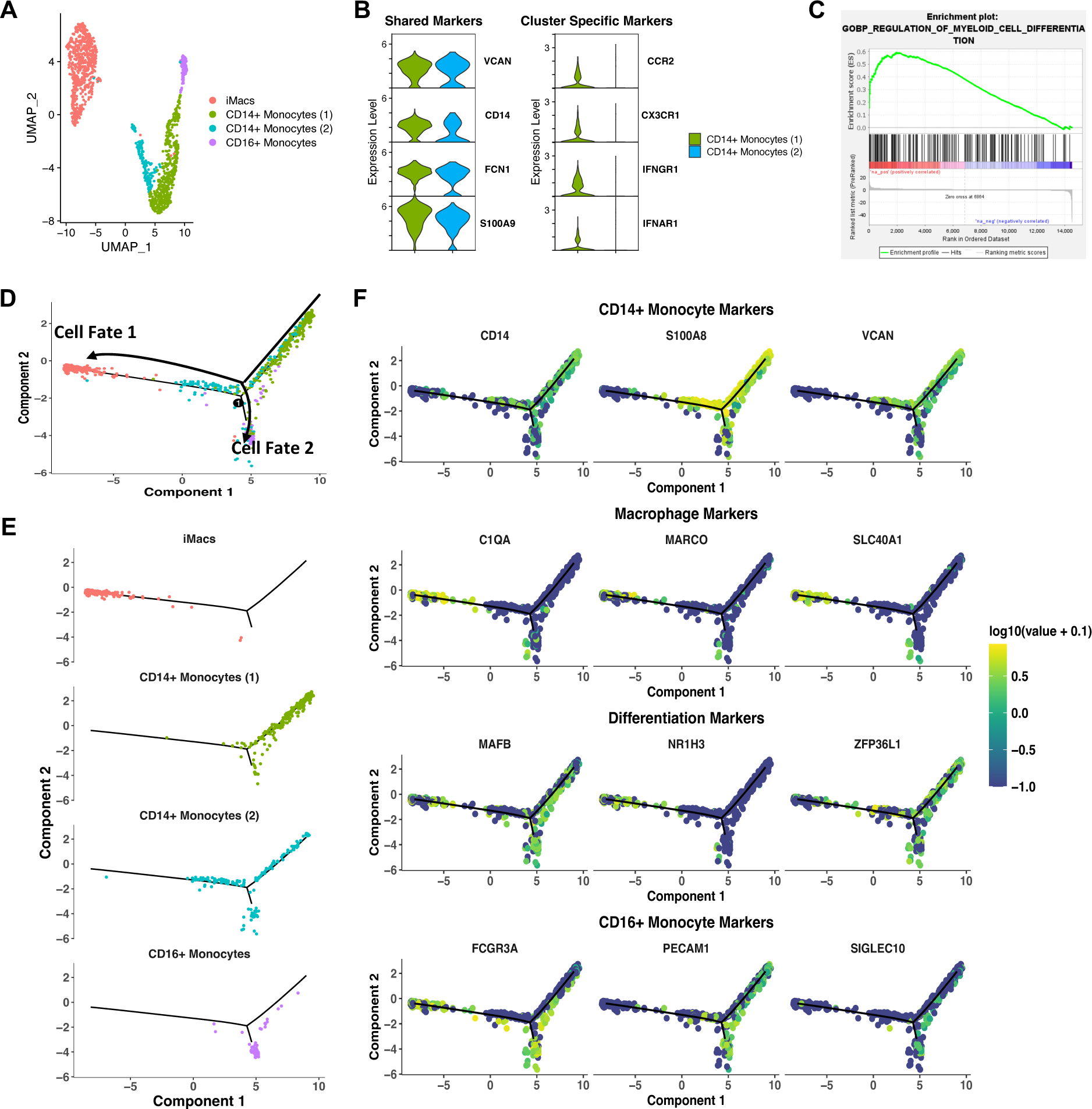
Analysis of monocyte/ macrophage differentiation trajectories during liver inflammation in CHB patients before antiviral treatment. **A)** iMacs, both CD14+ Monocytes and CD16+ Monocytes were reclustered using Seurat during liver inflammation using UMAP dimensionality reduction. **B)** Violin plots of CD14+ monocyte-defining genes, all selected genes have an adjusted p value <0.05. **C)** Enrichment plot from pathway analysis done on CD14+ Monocytes at baseline versus week24 of TAF treatment. Each vertical line represents a differentially expressed gene belonging to the pathway. **D)** Differentially expressed genes between clusters were used to generate hypothetical developmental relationships using the Monocle algorithm. **E)** Individual clusters along Monocle trajectory. **F)** Gene expression of CD14+ Monocyte-(first row), iMacs-(second row), Differentiation-(third row) and CD16+ Monocyte-(fourth row) defining genes along the pseudotime trajectory. iMacs, inflammatory macrophages.

To characterize developmental relationships between CD14+ monocytes, CD16+ monocytes, and the iMacs, Seurat-defined clusters were subclustered and superimposed on a pseudotime trajectory produced by the Monocle algorithm(*38*) (Fig. 4A and 4D). Monocle analysis revealed both CD14+ monocyte populations at different points in a successive manner along the same trajectory, representing cells in different states of differentiation, which matched the GSEA prediction (Fig. 4D). Furthermore, the trajectory suggests 2 potential cell fates, where cell fate 1 corresponds to the iMacs and cell fate 2 corresponds to CD16+ Monocytes. The majority of CD14+ Monocytes (2) oriented towards cell fate 1, implying the iMacs are derived from CD14+ Monocytes (Fig. 4E).

Gene expression was then plotted as a function of pseudotime to track changes along the trajectory during monocyte-to-MΦ differentiation. *CD14, S100A8* and *VCAN* expression was high in CD14+ Monocytes (1), lower in CD14+ Monocytes (2) and absent in the iMacs, suggesting the cells lose monocyte identity over the course of the trajectory. Loss of monocyte identity was associated with gain of *C1QA, MARCO* and *SLC40A1* in the iMacs (Fig. 4F). Monocyte markers of differentiation, *MAFB* and *ZFP36L1,* decreased along the path to iMacs, while the MΦ lineage marker *NR1H3* increased (Fig 4F). CD16+ monocytes progressively increased expression of *FCGR3A, PECAM1* and *SIGLEC10* and retained *MAFB* and *ZFP36L1,* suggesting they could also originate from the differentiation of CD14+ Monocytes (Fig. 4F). This analysis indicates a differentiation pathway from blood monocytes to a transitional CD14+ monocyte population before assuming the monocyte-derived MΦ phenotype within the liver.

### Tissue resident CD8 T cells and Kupffer cells contribute to the development of the inflammatory MΦs in the liver of CHB patients

To identify signals driving monocyte differentiation, the NicheNet algorithm(*39*) was used to infer potential ligand-receptor interactions in iMacs during inflammation. Among the top predicted ligands for iMac differentiation were Apolipoprotein E (ApoE), Interferon Beta (IFN-β), Gamma (IFN-γ), CXCL12 and colony stimulating factor 1 (CSF-1), also known as macrophage colony stimulating factor (M-CSF) (Fig. 5A–5C). ApoE was predicted to up-regulate *C1QA, C1QB,* and *HMOX1* expression. Type-I IFN was anticipated to upregulate interferon stimulated genes (ISGs) *IRF1* and *STAT1* while type-II IFN was predicted to upregulate *GBP1, HLA-DRA,* and *SOD2* expression (Fig. 5C).

**Figure 5.**
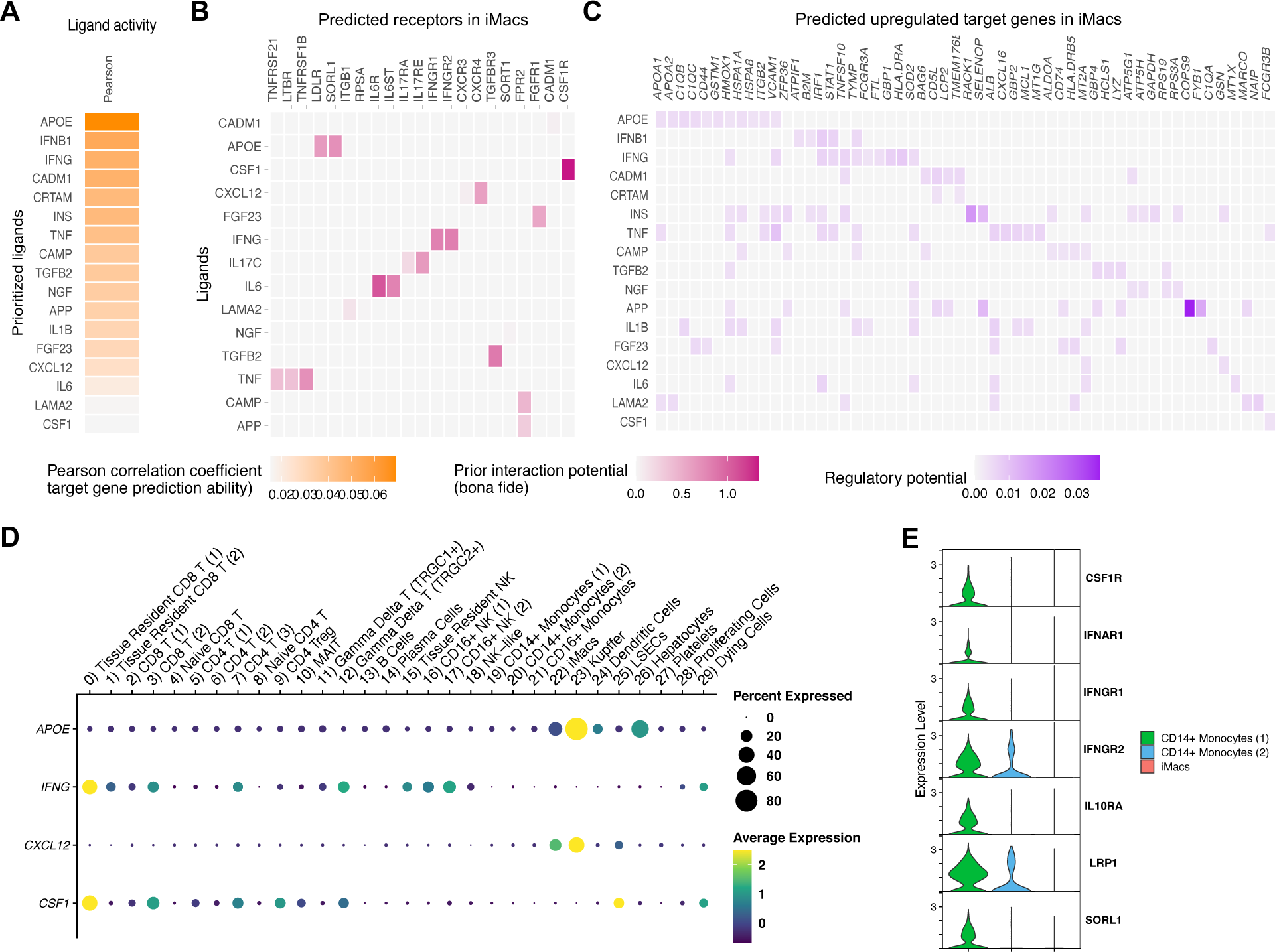
Predicting ligand-receptor interactions during liver inflammation suggests an inflammatory loop between the iMacs and CD8 T cells which will serve to guide in vitro monocyte-to-macrophage differentiation. **A, B, C)** Schematic representation of NicheNet analysis of ligand-receptor pairs inducing the differentially expressed gene profile of the iMacs during liver inflammation (baseline). **A)** Potential ligands, **B)** receptors, and **C)** target genes that may be driving Mac differentiation and activation at baseline. Only ligand receptor interactions that have been previously reported included in literature are included in **C. D)** Dot plot of NicheNet analysis predicted ligands for all Seurat clusters. **E)** Violin plot of NicheNet analysis predicted receptors on the iMacs and both CD14+ Monocyte clusters, all selected genes have an adjusted p value <0.05. iMacs, inflammatory macrophages.

Next, we identified potential sources of these ligands within our dataset to predict intercellular crosstalk from the scRNAseq data. Two key ligands, IFN-γ and CSF-1, were mapped to tissue resident CD8 T cells (Cluster 0) whereas ApoE was highly expressed by KCs (cluster 23), which also highly expressed the chemokine CXCL12 (Fig. 5D). Type-I IFN transcripts were not detectable within the dataset. The predicted interaction between iMacs and tissue resident CD8 T cell was consistent with IMC data described above (Fig 2F and 2G). We also observed a progressive decrease in the receptors for these ligands as monocytes progressed through differentiation from monocyte (1) to monocyte (2) to iMacs (Fig. 5E). Therefore, local signals from the inflamed liver microenvironment are responsible for differentiating the recruited monocytes into the iMacs.

### Predicted ligands drive iMac differentiation from blood monocytes

Our analysis predicted iMacs differentiate from monocytes. Therefore, we evaluated whether we could generate the iMac phenotype from monocytes *in vitro*. (Fig 6A). M-CSF was used to prime monocytes for differentiation and then individual cytokines with the highest predictive potential were used to induce markers corresponding to the transcriptional phenotypes: ISGs (*IFI27* and *IFIT3*)(*35, 36*), markers of inflammation (IL-18)(*32*), *FCGR3A* (CD16), and recent monocyte-to-MΦ differentiation (*NR1H3, ZFP36L1* and *MAFB*)(*4, 31, 40*). Loss of *VCAN expression* and increased *C1QA expression* were used to validate the transition from monocyte to MΦ(*24*)*. CETP* was used as a negative control to exclude KCs.

**Figure 6.**
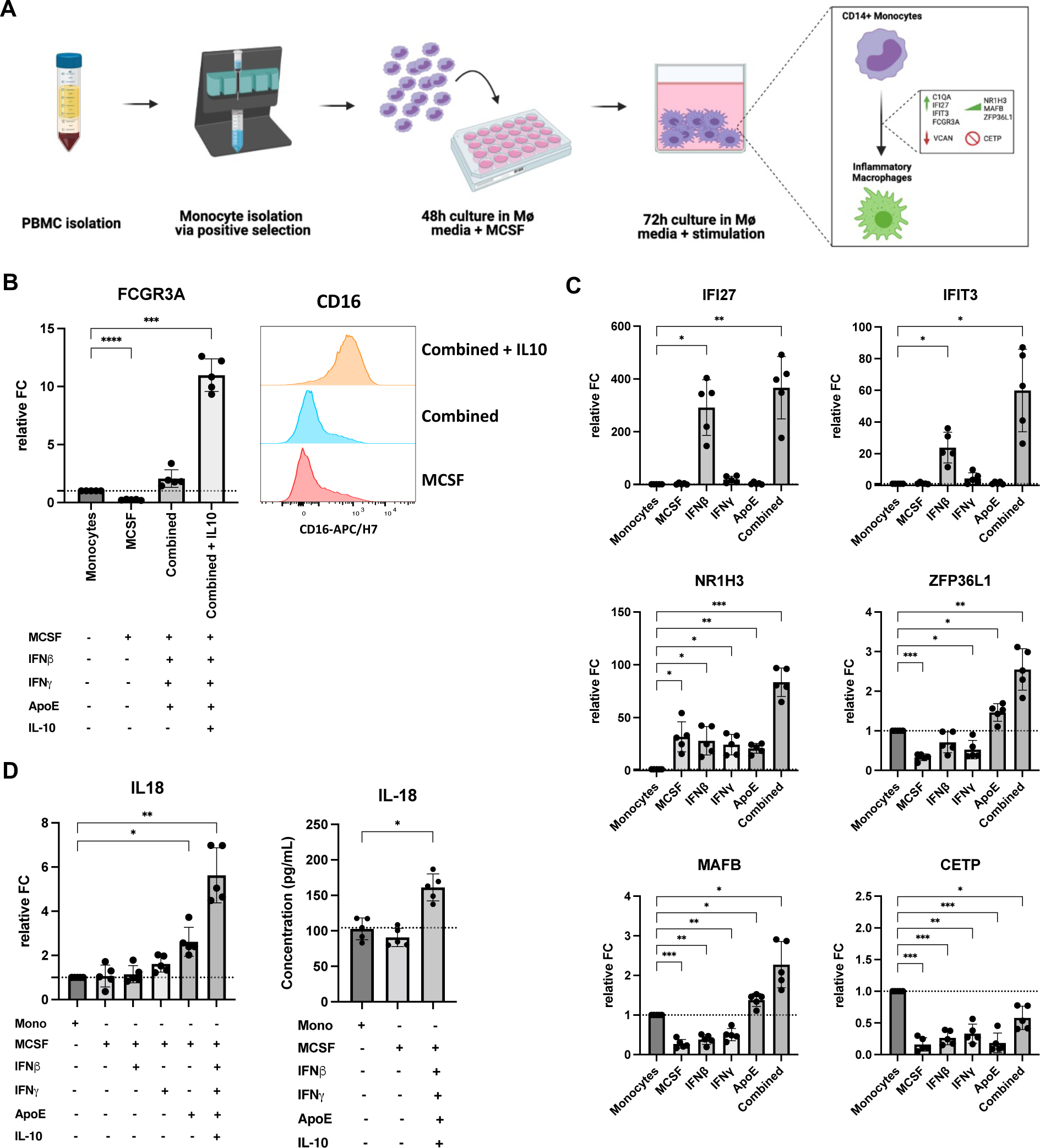
*In vitro* monocyte-to-macrophage differentiation suggests type I and II IFN stimulation upregulates key markers from inflammatory macrophages predicted by scRNAseq analysis. **A)** Workflow used for *in vitro* monocyte-to-macrophage differentiation using NicheNet ligands with predicted phenotype. **B)** Expression of CD16 (FCGR3A) at transcriptional level (left) and protein level (right) using IL10 stimulation in addition to NicheNet predicted ligands. CD16 protein expression measured from MFI. **C)** Real-time qPCR analyses of relative fold change for mRNA expression on differentiated Macs using individual and combined ligands predicted by NicheNet. **D)** Expression of IL-18 at transcriptional-(top row) and protein-level (bottom row) using NicheNet ligands. *P* values determined by repeated-measures one-way ANOVA, (\**P* <0.05, \*\**P* <0.005, \*\*\**P* <0.001). Results showed are representative of **n=**5 experiments. Mono, Monocytes; Macs, macrophages; MCSF, macrophage colony stimulating factor; IFN, interferon; MFI, median fluorescence intensity. “Combined” ligands include MCSF, IFN-β, IFN-γ and Apolipoprotein E (ApoE).

Exploratory *in vitro* experiments revealed CD16 could not be induced by any of the predicted ligands (Fig. 6B and Suppl. Fig. 3A). For that purpose, we returned to NicheNet to predict ligands that specifically upregulate CD16. IL-10 had the highest regulatory potential score (Suppl. Fig. 3B). *In vitro* monocyte-to-MΦ differentiation validated IL-10’s capacity to upregulate CD16 at the transcriptional (Fig. 6B, left column) and protein level (Fig. 6B, right column) in the presence of the other predicted cytokines under the condition “combined”, which includes M-CSF, IFN-β, IFN-γ and ApoE. ISGs *IFI27* and *IFIT3*, were induced by IFN-β. *NR1H3* was induced by M-CSF while ApoE significantly increased expression of *ZFP36L1* and *MAFB* (Fig. 6C). The combination of all the predicted cytokines had a synergistic effect on the upregulation of each of the predicted genes (Fig. 6C and Suppl. Fig. 3C). *CETP* was not induced under any condition and M-CSF decreased *VCAN* and increased *C1QA* (Fig. 6C; Suppl. Fig. 3A).

Once the differentiation conditions for iMacs were defined, their capacity to induce inflammation was assessed. IL-18 plays a key role in tissue damage during infections and chronic inflammatory diseases(*41*). IL-18 was significantly induced at the transcriptional level by ApoE alone and further enhanced by the combined cytokine cocktail (Fig. 6D, left). Consistent with transcriptional activation, we also measured increased IL-18 protein in cell culture supernatant (Fig. 6D, right). Further evaluation revealed that the differentiated iMacs represent a unique state between M1 and M2 *in vitro* differentiated MΦ. They express M1 markers, such as HLA-DR and CD40, but lack CD86 and express the M2 marker CD16, but lack CD206 and CD209 (Suppl. Fig. 4A). At the functional level, iMacs were largely inflammatory by their release of cytokines such as IL-1β, IFN-α*2,* MCP1, IL-, IL-18 and IL-23, but also secrete IL-10. iMacs differed functionally from M1 MΦs by reduced production of TNF-α, IL-6 and IL-12p70 (Suppl. Fig. 4B). Therefore, we could validate the differentiation pathway from monocytes to iMacs predicted by the ligand receptor analysis. The iMacs displayed a phenotypic profile sharing characteristics with both M1 and M2 MΦs but were highly inflammatory based on their cytokine production profile.

### The inflammatory MΦs are long lived but remain quiescent after resolution of inflammation

Animal models indicate that embryonic KCs are replaced by monocyte-derived MΦs after liver damage. As chronic hepatitis is a driver of cirrhosis, we wanted to understand if the iMacs were short-lived or if they remained in the liver where they could contribute to future inflammatory events. To address this question, we incorporated an additional scRNAseq dataset from CHB patients. These patients had active hepatitis then received 4 years of nucleos(t)ide analogue (Nuc) therapy to reduce HBV replication to undetectable levels and normalize ALT levels(*42*). The data from this analysis were integrated with data from Figure 3 (HBV-inflamed, uninfected healthy(*25*) and cirrhotic(*13*) human livers) and the same 5 MΦ populations were identified (Fig 7A and Fig 3D). Again, iMacs were enriched in CHB patients compared to healthy and cirrhotic livers, but it was visually apparent that iMacs from patients with active disease occupied a distinct space within the cluster compared to Nuc-treated patients (Fig 7B). iMacs from the inflamed and Nuc-treated patients displayed a similar transcriptional profile: *SLC40A1, MARCO, IFI27, SPI1* (PU.1), *FCGR3A* and the absence of *CD14* (Fig 7C, first column). However, the markers of immune activation (IL18 and LGALS9)(*32, 33*), IFN signaling (IFIT3 and GBP5)(*35, 36*), and recent monocyte-to-MΦ differentiation (MAFB and ZFP36L1)(*30, 40, 43*) were absent in the patients on Nuc therapy (Fig 7C, second column). This supports the concept that iMacs populate the liver during inflammation, but once it resolves, they remain in a long-lived quiescent state.

**Figure 7.**
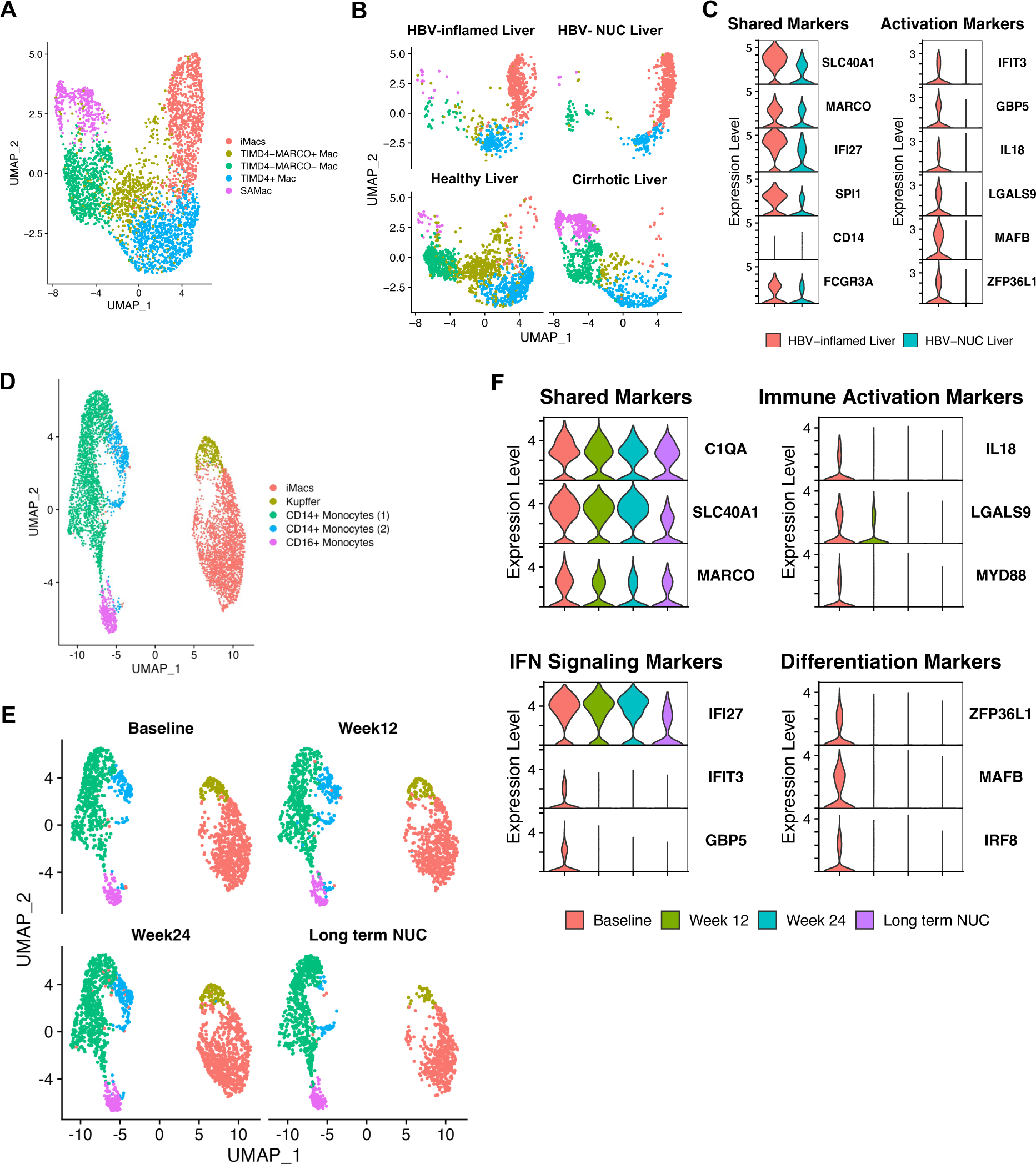
Comparison of iMacs during HBV reactivation to healthy, cirrhotic and CHB patients on long term antiviral human livers at the single cell level. **A)** Clustering of Macs from healthy (**n=**5), cirrhotic (**n=**5), HBV-inflamed (**n=**5) and long term NUC (**n=**5) human livers using UMAP dimensionality reduction. **B)** UMAP dimensionality reduction of liver Macs by liver condition. **C)** Comparison of cluster defining genes for iMacs in violin plot during liver inflammation and in the absence of liver inflammation. **D)** Clustering of CD68+ cells during different stages of CHB using UMAP dimensionality reduction. **E)** UMAP dimensionality reduction of CD68+ cells by stage of CHB. **F)** Violin plots of iMacs-defining genes by stage of CHB. All selected genes have an adjusted p value <0.05. iMacs, inflammatory macrophages; Macs, macrophages; SAMacs, Scar-associated macrophages; NUC, nucleoside analogue.

Finally, we compared the distribution of precursor monocyte populations and expression of key genes in relation to time on Nuc therapy to better understand how these change as liver damage resolved. Myeloid characterization yielded the same MΦ and monocyte clusters as previously defined (Fig 7D). Interestingly, CD14+ monocyte (2), which represented the transitional monocyte population in our Monocle analysis (Fig 2D), was almost undetectable in long-term Nuc-treated patients (Fig 7E). We then compared changes in gene expression over time. While phenotypic markers (*C1QA, SLC40A1, MARCO*) were stable over time, markers of immune activation, IFN signaling, and recent monocyte-to-MΦ differentiation were decreased by 12w of antiviral therapy and remained undetectable at 24w and in long-term Nuc treated patients (Fig 7F). Taken together, these data support a model where liver inflammation gives rise to a unique inflammatory MΦ that is capable of establishing a long-term survival niche in the human liver, poised to contribute to progressive liver damage in the future.

## Discussion

Liver inflammation puts 800 million people at risk for cirrhosis (*9*), 290 million of which have chronic hepatitis B (*44*), with MΦs acting as key regulators of inflammation. While we anticipated MΦ activation from our previous analysis of serum markers in patients with hepatitis (*20*), our data show the implications of MΦ activation are not transient. Monocytes infiltrating the inflamed human liver integrated complex environmental signals to establish a long-lived monocyte-derived MΦ population biased towards inflammation. Imprinting a long-lived MΦ population whose abundance positively correlated with CD8 T cells responsible for liver damage suggests they could represent the fulcrum, tilting the scale towards progressive tissue damage during chronic liver disease.

The concept of trained immunity has been established for innate immune cells that lack cognate antigen-specific receptors (*45*). Exposure to a pathogen imprints a functional response based on the inflammatory environment that promotes a pre-defined response upon re-exposure. Despite monocytes being short-lived, both BCG vaccination and *Candida albicans* infection can induce a state of trained immunity in adults (*46, 47*). We previously demonstrated that trained immunity occurs at birth in children born to HBV+ mothers, with a CD14+ monocytes biased toward a Th1-mediated response (*41,44*). *In vitro*, monocytes exposed to inflammatory stimuli adapt their transcriptional profile to environmental signals to influence MΦ differentiation, opening a pathway to reshape the liver MΦ landscape (*48*). Our longitudinal data show this happens over extended time in patients, with the transitional CD14+ monocyte (2) population still present after 6m of therapy, but absent after long-term antiviral treatment. Disappearance of the transitional monocyte (2) population, and stability of the iMac pool after long-term therapy, suggests iMacs are not constantly being repopulated from blood and suggests this recruitment occurs upon niche availability caused by KC death. The iMac population may establish the capacity for self-renewal, similar to what was observed after Leishmania infection (*49*).

The imprinted transcriptional profile of iMacs was clearly distinct from KCs, which occupied the same inflamed organ. KCs are biased towards an anti-inflammatory profile and remove dying cells to prevent local inflammation (*2*). However, this can limit their ability to become inflammatory and can potentially be exploited by pathogens. Instead, recently recruited monocyte-derived MΦs are poised to become inflammatory and display enhanced antimicrobial capacity(*50*). However, their fate upon resolution of inflammation, cell death or engraftment into the tissue, has not been well defined. Our data from CHB patients after long-term antiviral therapy indicate that iMacs use the niche created in the liver during hepatitis to establish a long-lived population, which may posses a lower threshold for future activation.

When iMacs were further compared to MΦs from the healthy and cirrhotic livers, the distinction remained. The relatively fewer macrophages in the KC cluster were dispersed across previously identified macrophage phenotypes in healthy and cirrhotic datasets, while iMacs remained separate and highly enriched in the inflamed liver. The distinction extended to spatial localization, with iMacs clustering around the portal areas while KCs were dispersed throughout the liver lobules. These data point towards a concept of hyper-local regulation by distinct MΦ subsets across the liver lobules. Whether this spatial distribution endures in patients on long-term antiviral therapy remains to be determined but knowledge of localization and cell-cell interactions could yield strategies to suppress liver damage across different etiologies of chronic liver disease.

The functional profile of iMacs, their dependence on CD8 T cell-derived cytokines, and co-localization with CD8 T cells, suggests that they operate in an inflammatory loop. IFN-γ and M-CSF were essential factors to induce the iMac phenotype. The iMac phenotype, being similar to M1-like MΦs, was associated with inflammatory cytokine production, including production of IL-18. IL-18, in combination with IL-12, activates CD8 T cells to express IFN-γ and FasL in an antigen-independent manner (*51*). Consistent with these data, we recently demonstrated that tissue-resident CXCR6+ CD8 T cells respond to these cytokines to drive antigen-independent hepatocyte killing through FasL(*21*). This creates a positive feedback loop where IL-18 stimulation can lead to the release of IFN-γ which in turn regulates IL-18 release from the iMacs (*32*). This cycle is broken by the introduction of antiviral therapy, which rapidly suppresses viral replication.

In addition to tissue-resident CD8 T cells, KCs appear to play a central role in orchestrating iMac differentiation. Our IMC data confirmed that KCs are not completely eliminated and remain in the inflamed liver. However, during inflammation, they remained distributed throughout the lobule, while iMacs were present mainly in portal regions. KCs expressed the highest level of CXCL-12 and may be responsible for driving CXCR4+ monocytes to the liver. ApoE is a physiological regulator of lipid homeostasis with the capacity to modulate the immune response against pathogens, such as malaria, mycobacteria, and viruses (*52*). Its role in monocyte-to-MΦ differentiation was that of an enhancer, inducing *MAFB* and *ZFP36L1* to drive monocyte to MΦ transition and work in concert with IFN-γ to enhance IL-18 expression. When put into context, KC activation through danger signals associated with HBV replication could trigger the recruitment and activation of monocytes. Infiltrating monocytes integrate inflammatory signals from KCs and CD8 T cells, synergistically increasing their inflammatory transcriptional signature to establish a phenotype that propagates liver damage (*10, 46-47*).

We knew that monocyte-derived MΦs were present in the human liver during inflammatory events. However, previous human studies captured a snapshot of the liver environment during inflammatory disease because core biopsies are collected to confirm diagnosis or staging, not dynamic changes in activation profiles. Therefore, previous studies lacked dynamic longitudinal data demonstrating the transition from inflammatory monocyte-derived MΦ during liver damage to a long-lived MΦ poised for reactivation. With this in mind, the role of iMacs in the different stages of CHB, which are defined by different degrees of viral replication and liver damage, require further investigation. We hypothesize that iMacs arise as CHB patients transition from an immunotolerant state of high viral load with normal ALT levels to an immune active state where ALT begins to rise. The iMacs become established in the liver and likely contribute to inflammation associated with HBV reactivation upon therapy withdrawal, where we have observed a MΦ signature(*23*). Understanding how the iMac population is established and how it contributes to the inflammatory loop propagating liver damage could provide key insights to manage liver inflammation across etiologies to slow the progression of chronic liver diseases.

