## Supplementary Material for "Virus-associated Inflammation Imprints an Inflammatory Profile on Long-lived Monocyte-derived Macrophages in the Human Liver"

### Methods

#### Lead contact

#### Data availability

The datasets generated during the current study are available from the corresponding author on reasonable request. The datasets are available in the National Center for Biotechnology Information Gene Expression Omnibus (GEO) repository: GSE216314.

#### Code availability

R scripts are available from the corresponding author on reasonable request. Any additional information required to reanalyze the data reported in this paper is available from the lead contact upon request.

#### Experimental models and subjects' details

Chronic hepatitis B patients

11 patients were included in this study to analyze blood and liver FNAs (NCT: NCT04070079). Inclusion criteria were chronic hepatitis B (HBsAg (+) ≥ 6 months); age >18 years; elevated ALT levels, defined as >19 IU/l for females and >30 IU/l for males

(with ULN defined as >25 IU/l for females and >35 IU/l for males); HBV DNA >10000 IU/ml for HBeAg (+) and >1000 IU/ml for HBeAg (-) patients; adequate contraception. An overview of baseline characteristic is given in suppl. table 1. Exclusion criteria were antiviral or IFN treatment in the previous 6 months; immunosuppressive treatment in the previous 6 months; treatment with an investigational drug in the previous 3 months; history of decompensated liver cirrhosis; liver transplantation; co-infection with HCV, HDV, or HIV; other significant liver disease (such as alcoholic or drug-related liver disease, autoimmune hepatitis, hemochromatosis, Wilson's disease or α1 antitrypsin deficiency); estimated glomerular filtration <50 ml/min/1.73m$^2$ or significant renal disease; α-fetoprotein >50 ng/ml; pregnancy or breast feeding; other significant medical illness that might interfere with the study (e.g. immunodeficiency syndromes or malignancies); substance abuse. Of the 5 patients analyzed by scRNAseq, 4 were male and one was female. Two patients were HBeAg (+), and three patients were HBeAg (-). Due to low numbers, no analysis of sex influence was included[41].

Human donors of peripheral blood mononuclear cells (PBMCs)

5 healthy human donors were included to obtain PBMCs from whole blood. Mean age at the time of donation was 32.1 years (range 23–53 years). 3 patients were male, and 2 patients were female. Due to low numbers, no analysis of sex influence was included. Informed consent was obtained from all subjects.

**Methods details**

### Ethics statement

This investigator-initiated clinical study (NCT: NCT04070079) was approved by the University Health Network Research Ethics Board (CAPCR ID: 18-5748). All patients provided written informed consent.

### Study design

This was an investigator-initiated, open-label phase 4 study at the Toronto Centre for Liver Disease, Canada. Chronic hepatitis B patients with  viral load HBV DNA> 2000 IU/mL and liver damage, measured by elevated levels of serum alanine aminotransferase (ALT) represented as fold increase over normal values (* upper limit of normal (*ULN >1)) started therapy with 25 mg daily of tenofovir alafenamide (TAF) for a duration of 48 weeks and were offered to continue therapy after the end of the study. Blood and FNA samples were collected at baseline, week 12 and week 24. Additional blood samples were collected at week 36 and 48.

### Analysis of blood markers of HBV infection

Markers of HBV infection in patients' blood were measured at baseline, and at week 12 and week 24 after starting antiviral therapy. HBV DNA levels and alanine aminotransferase (ALT) times upper limit of normal (ULN) were measured using AmpliPrep TaqMan (Roche) and Advia (Siemens), respectively. HBsAg and HBeAg were measured using the Architect assay (Abbott). Measurement of blood markers of HBV infection was done by the Laboratory Medicine Program of Toronto General Hospital/University Health Network. The clinical characteristics of the patients' profiles is listed in Supplementary table 1.

Liver FNA collection

Liver FNAs were collected by a hepatologist. The site of liver puncture for FNA was determined by bedside ultrasound and patients underwent local anesthesia with 2% lidocaine. 22- or 25-gauge needles were used for aspiration of cells as previously described (https://journals.lww.com/hep/Abstract/9900/Single_cell_RNA_sequencing_of_liver_fine_needle.421.aspx). 20,000 cells per sample were subjected to scRNAseq.

Determination of cytokine concentrations from plasma

Detection of soluble CD163 (sCD163), Interleukin (IL)-18 and Galectin-9 from plasma collected from chronic hepatitis B patients were performed using a custom made Human Magnetic Luminex Assay (R&D Systems) at baseline, and 12 and 24 weeks after starting antiviral therapy. The protocol was performed according to the manufacturer's instructions. Data were acquired using a MAGPIX instrument (Luminex) and concentrations were calculated using xPONENT software (version 4.2).

Tissue Staining for Imaging Mass Cytometry

Tissue staining by imaging mass cytometry and image analyses were conducted as previously described (https://insight.jci.org/articles/view/146883), using paraffin-embedded formalin fixed liver tissue slides from chronically HBV-infected patients (28 immune active and 6 immune tolerant subjects).

#### Gating strategies for FCS files for CD8 T-cells, iMacs and Kupffer Cells

CSV files exported from R were converted into flow cytometry standard (fcs) files, and ± cutoff values for each marker and sample were manually chosen in FlowJo based on biaxial plots using the CD45 channel for comparison. These cutoff values were then imported into R and used for gating, employing the flowCore and flowWorkspace packages, and subsequent statistical analyses. CD8 T cells were further gated for $CD3^+CD8^+$ while myeloid cells were gated for $CD68^+$, $CD16^+$, and $CD14^+$ cells. Notably, expression patterns for CD68, CD16, and CD14 were highly colocalized with similar characteristics in quantitative and phenotype analyses. Therefore, we simplified our language and data presentation by using CD68 expression to define hepatic macrophages—defining iMacs as CD68+CD16+CD14- CD45+ cells and Kupffer cells as CD68+CD16+CD14+CD45+ cells. Gated immune subsets were then analyzed for phenotype marker expression. Results were normalized as hepatic cellular density in counts per $mm^2$ ROI and as percentages for each cell subset of interest.

#### 10x Genomics sample processing for single-cell RNA sequencing

For scRNAseq, samples were prepared as outlined by the 10x Genomics Single Cell 5' Reagent Kit user guide with a maximum capture target of 3,000 cells. Briefly, after droplet generation, samples were transferred onto a pre-chilled 96-well plate, heat sealed, and cDNA was generated overnight. The next day, cDNA was recovered using Recovery agent (10x Genomics), and then it was purified using a Silane DynaBead mix (ThermoFisher) following manufacturer instructions. Purified cDNA was amplified for 14 cycles, then it was re-purified using SPRIselect beads (Beckman Coulter). cDNA concentration was measured using a Bioanalyzer (Agilent Technologies). 5' cDNA libraries were prepared as outlined by the 10x Genomics' Single Cell 5' Reagent Kit user guide, with modifications to the PCR cycles based on the calculated cDNA library input. Sequencing libraries were generated with unique sample indices for each sample and quantified.

The molarity of each library was calculated based on library size as measured by the Bioanalyzer (Agilent Technologies) and qPCR amplification data. Samples were pooled and adjusted to 10 nM, then diluted to 2 nM. Each 2 nM pool was denatured using 0.1 NaOH at equal volumes for 5 minutes at room temperature. Library pools were further diluted to a final loading concentration of 14 pM. 150 μl were loaded into each well of an 8-well strip tube and loaded onto a cBot (Illumina) for cluster generation. Samples were sequenced on the HiSeq 2500 (Illumina) system.

Raw sequencing data were aligned to the human genome reference sequence GRCh38 combined with HBV genome and convered to unique molecular identifier (UMI) counts per gene per cell using the CellRanger (10x Genomics) analysis pipeline.

Cell clustering, differential expression, and pathway analysis

The CellRanger-processed filtered feature matrices were analyzed using the Seurat R package (version 3.2.3)[62]. The raw digital gene expression matrix (UMI counts per gene per cell) from each sample was filtered, normalized, and clustered. To preserve high quality cells, filtering was performed as follows: Cells with low transcript counts (<200 UMIs) and high mitochondrial transcript ratio (>30%) were removed. Genes that appeared in less than 3 cells were removed as well. Normalization was performed using the scran R package (version 1.18.5)[63]. After normalization, data from all samples was integrated and scaled using Seurat. Clustering was performed using standard Seurat package procedures. Principal component analysis (PCA) was used to reduce the number of dimensions representing each cell. The number of components used was determined based on the elbow of a scree plot. A shared nearest neighbor graph was built from distances computed in principal component space with k.param set to 20 PCA dimensions as inputs. Selection of a biologically relevant number of clusters was based on differential expression between neighboring clusters which were identified as the next-nearest cluster to each cell after the cell's assigned cluster. Clusters were visualized using uniform manifold approximation and projection (UMAP) coordinates of the principal components as implemented in Seurat. Clusters were identified using the Louvain algorithm with a resolution set to 0.9. Cell-type identities for each cluster were determined manually based on canonical transcriptional markers of well-defined parenchymal/ non-parenchymal liver cells. Differential gene expression analysis was performed using the standard area under the curve (AUC) classifier to assess significance. We retained only those genes with a log-transformed fold change of at least 0.15 and expression in at least

25% of cells in the cluster under comparison (Supplementary Table 2). Further clustering was performed in CD68+ clusters to reveal specific myeloid populations. At each stage of this process, data was scaled and PCA was performed to reveal biologically relevant clusters. We removed clusters expressing more than one unique lineage signature in more than 25% of their cells from the dataset as probable doublets.

For the comparison of liver MΦs between different stages of liver disease and healthy human livers described in Figures 3 and 7, scRNAseq datasets were obtained from publicly available reference datasets. Further information on GEO accession number for the datasets is listed in the key resources table. For downstream analysis, individual samples were filtered and normalized using the same parameters previously described. The data was integrated using Seurat, then further clustering was performed on liver MΦs and differential gene expression was determined (Supplementary table 3). Downstream analysis following data integration was performed as previously described.

Pathways enriched in specific clusters in Fig 2D and 2E were elucidated using gene set enrichment analysis (GSEA)[40]. Briefly, gene rank lists were compiled in Seurat, and GSEA was performed using the fgsea R package (version1.16.0). c5.go.bp.v2022.1.Hs.symbols.gmt from [http://www.gsea-msigdb.org/gsea/msigdb/collections.jsp] was used to identify enriched cellular pathways in GSEA analysis.

All heatmaps, UMAP visualizations, violin plots, dot plots and feature plots were produced using Seurat functions in conjunction with the ggplot2 and pheatmap R packages. Information on the respective vignettes is provided in the "additional resources" section.

Pseudotime trajectory analysis

To generate cellular trajectories to infer developmental relationships between Monocytes and the iMacs at the time of liver inflammation we used the monocle R package (v2.18.0)[42]. We ordered cells in a semi-supervised manner based on their Seurat clustering and using the top 2,000 highly variable genes as input we sorted cells in pseudotime. Dimensional reduction was performed using the function “DDRTree” to infer potential developmental paths and pseudotime ordering was performed using the “orderCells” function using CD14+ Monocytes (1) as the start point of the trajectory. Differentially expressed genes along this trajectory were identified using generalized linear models via the ‘differentialGeneTest’ function in monocle. The remaining parameters were default.

Ligand-receptor interaction analysis

To understand cell-cell interactions at the time of liver inflammation and potential upstream signals driving monocyte differentiation into the iMacs we used the nicheNetr R package (v.1.0.0)[43]. Ligand-target prior model, ligand-receptor network, and weighted integrated networks were imported from NicheNet data sets. iMacs were set as receiver/target cell population and all liver FNA clusters were set as potential senders. Baseline was set as the condition at which receiver cells were affected by other cells and week 24 was set as the steady-state condition. Then, potential ligands were ranked based on the presence of their target genes in the gene set of interest. The top 20 ligands and their cognate target genes were inferred and visualized in a heatmap and validated on dot plots. Genes expressed in at least 10% of the cells in one cluster were considered expressed in this cluster. The remaining parameters were default.

Monocyte isolation from human blood

Whole blood was collected from healthy human volunteers into vacutainer tubes containing the anticoagulant ACD (ThermoFisher). Total blood was diluted in a 1:2 ratio with 2% Knockout Serum Replacement (ThermoFisher). Diluted blood was layered over Lymphoprep solution (StemCell) in SepMate-50 tubes (StemCell). Density centrifugation was used to remove red blood cells and the supernatant containing PBMCs was transferred and were washed 2x with PBS (ThermoFisher) + 2% Knockout Serum Replacement (ThermoFisher) to remove remaining Lymphoprep solution and platelets. PBMCs were counted.

CD14+ Monocytes were purified by positive selection using human CD14+ microbeads (Miltenyl Biotec) following manufacturer's instructions. Purity of CD14+ monocytes was assessed and was found to be ≥ 85%. Pure CD14+ Monocytes were resuspended in MΦ media which included RPMI (ThermoFisher) supplemented with 10% human serum (Sigma-Aldrich), 1% penicillin/streptomycin (Lifetech) and 1% GlutaMAX (ThermoFisher).

*In vitro* generation of monocyte-derived MΦs

CD14+ Monocytes were plated at a seeding density of 1 x$10^6$ cells/mL in MΦ media on 48-well untreated plated (ThermoFisher). A parallel incubation was done on 96-well untreated plates (ThermoFisher) for cytokine release measurement upon TLR agonist stimulation. To trigger monocyte differentiation into iMacs, cells were plated at 37°C in a humidified incubator with 5% $CO_2$ in the presence M-CSF (Goldbio) at 100 ng/mL for 48 hours. Cells were then polarized for additional 72 hours with cytokines predicted by the NicheNet algorithm. Indicated cytokines were used in the following concentrations: IFN-β (Goldbio) at 100 IU/mL, IFN-γ (Goldbio) at 100 IU/mL, ApoE (Sigma) at 5 µg/mL, and IL-10 at 5 ng/mL (Goldbio). Additional cytokines were used for comparison of inflammatory MΦ against canonical M1- (pro-inflammatory) and M2- (anti-inflammatory) MΦ using: IFN-γ (Goldbio) at 100 IU/mL and LPS (Invivogen) at 1 µg/mL for M1 differentiation and IL-10 (Goldbio) at 25 ng/mL and IL-4 (GoldBio) at 25 ng/mL for M2 differentiation after 48 hours in the presence M-CSF (Goldbio) at 100 ng/mL. On day 5, supernatants were collected from the 48-well plates for measurement of cytokine release. This was followed by detachment of adherent MΦs using 10nM EDTA (Sigma-Alrich) in PBS at room temperature for flow cytometry analysis, or for lysis of adherent MΦ for RNA isolation. Further information on reagents used is listed in the key resources table.

RNA isolation and reverse transcription polymerase chain reaction (RT-PCR, qPCR)

Gene transcript levels were determined by qPCR. MΦ total RNA was extracted using the RNeasy Plus Mini Kit (Qiagen), resuspend in 14uL of ultrapure water and quantified using a NanoDrop 2000 (ThermoFisher), which was then converted to cDNA by reverse transcription using the High-Capacity cDNA Reverse Transcription Kit (ThermoFisher).

Both protocols were performed following the manufacturer's instructions. Gene expression was quantified with pre-designed TaqMan Fast Advanced Master Mix (Applied Biosystems) on a QuantStudio 6 thermocycler (Applied Biosystems) to investigate markers of iMacs and monocyte-to-MΦ differentiation. Gene expression was then normalized using the 2-ΔΔCq method relative to the expression of the housekeeping gene, GAPDH. Further information on reagents and primers used is listed in the key resources table.

Flow cytometry analysis of PBMC-derived MΦs

For flow cytometry analysis, fixable viability dye eFluor 450 (eBiosciences), 0.05:100 in PBS, was used for staining of dead cells for 10 minutes at room temperature. Subsequently, cells were washed and stained with extracellular antibodies for 30 minutes at 4°C: HLA-DR_BV605, 2.5:100; CD86_BV650, 1:100; CD40_BV711, 2.5:100; CD206_APC, 5:100; CD209_FITC, 2.5:100; and CD16_APC/H7, 2.5:100. All antibodies were diluted in staining buffer (PBS (ThermoFisher) + 1% BSA (Multicell) + 0.1% sodium azide (Sigma-Aldrich). Cells were washed and permeabilized using Cytofix/Cytoperm (BD Biosciences) for 15 minutes at 4°C and this was followed by staining with intracellular antibodies for 30 minutes at 4°C: CD68_PE/Cy7, 1:100, which was diluted in PermWash buffer: PBS (ThermoFisher) + 1%BSA (Multicell) + 0.1% sodium azide (Sigma-Aldrich) + 0.1% saponin (Sigma-Aldrich). Further information on antibodies used is listed in the key resources table. Cells were washed 2x, fixed and stored in PBS + 1% Paraformaldehyde (Sigma-Aldrich). The BD FACSSymphony A3 (BD Biosciences) cytometer was used. Data was analyzed with FlowJo (version 10.8.1).

Cytokine release assay

Cytokine protein levels secreted in the supernatant by differentiated MΦs were quantified using the LEGENDplex™ human inflammation panel 1 (BioLegend) for all markers, which included: IL-1β, IFN-*α*2, tumor necrosis factor (TNF)-*α*, MCP-1, Interleukin (IL)-6, IL-8, IL-10, IL-12p70, IL-18, and IL-33. All the experiments were performed following the manufacturer's instructions. The analytes were diluted in a 1:2 ratio before mixing with the samples. The samples were read using the BD FACSSymphony A3 (BD Biosciences) cytometer, and the data were analyzed using the cloud-based LEGENDplex™ Data Analysis Software (BioLegend). Further information on reagents used is listed in the key resources table.

**Statistical analysis**

All data is presented as the result of five or six independent experiments and expressed as mean +/- SEM. GraphPad Prism 9.3.0 (GraphPad Software, San Diego, CA, USA) and R version 4.0.3 was used to evaluate these data. The statistical analysis performed for each experiment is included in the figure legend. $P < 0.05$ was defined as statistically significant. Outlier analysis was performed using the ROUT method (Q = 1%). Number of replicates are indicated in the respective results sections and figure legends.

**Additional resources**

Seurat vignette : https://satijalab.org/seurat/articles/get_started.html

scran vignette: https://rdrr.igo/bioc/scran/f/vignettes/scran.Rmd

fgsea vignette: https://bioconductor.org/packages/release/bioc/html/fgsea.html

Monocle vignette : http://cole-trapnell-lab.github.io/monocle-release/docs/

NicheNet vignette : : https://rdrr.io/github/saeyslab/nichenetr/f/vignettes/seurat_wrapper.md


### Acknowledgments

We want to recognize the clinical support and administrative staff in the Toronto Centre for Liver Disease for their help in executing the study. Similarly, we thank the staff in the Princess Margaret Genomics Core facility and UHN-Sick Kids flow cytometry facility at University Health Network for their help in handling samples and data generation.


### Authors contributions

AJG, HLAJ and AG designed the study. HLAJ, SF, JJF enrolled patients for the study. AG, JJW, JJF designed experimental methods. J.D.S, DT, SK, SCK, DM, AM, DC, A.P, KMC conducted experiments or performed data analysis. J.D.S. and AJG wrote the main manuscript text and prepared the figures. All authors reviewed the manuscript.

### Declaration of interest

D.M. is employed by Fluidigm Inc. S.C.K. is employed by and stockholder of Gilead Sciences Inc. D.C. was formerly employed by Gilead Sciences Inc. and is currently employed by Bristol Myers Squibb. J.J.F. receives research funding by Abbvie, Arbutus Biopharma, Gilead Sciences Inc., Janssen Pharmaceuticals, Eiger Biopharmaceuticals, and Enanta Pharmaceuticals; and reports compensation from consulting/scientific advising for Abbvie, Arbutus Biopharma, Gilead Sciences Inc., and GlaxoSmithKline. S.F. receives research funding by Gilead Sciences Inc.; and reports compensation from consulting/scientific advising for Gilead Sciences Inc., Abbvie, Janssen Pharmaceuticals, Assembly Biosciences. J.J.W. is employed by and stockholder of Gilead Sciences Inc. A.G. was formerly employed by and stockholder of Gilead Sciences Inc. H.L.A.J. receives research funding by Abbvie, Gilead Sciences Inc., GlaxoSmithKline, Janssen Pharmaceuticals, Roche, Vir Biotechnology; and reports compensation from consulting/scientific advising for ALIGOS Therapeutics, Antios Therapeutics, Arbutus Biopharma, Eiger Biopharmaceuticals, Gilead Sciences Inc., GlaxoSmithKline, Janssen Pharmaceuticals, Merck, Roche, VBI Vaccines Inc., Vir Biotechnology, and Viroclinics Biosciences. A.J.G. receives research funding by Janssen Pharmaceuticals, GlaxoSmithKline, and Gilead Sciences Inc.; and reports compensation from consulting/scientific advising: Janssen Pharmaceuticals, Roche, GlaxoSmithKline, Vir Biotechnology, Finch Therapeutics, SQZ Biotech. The other authors declare no competing interests.

**Funding Sources**

A.G. was funded by the Canadian Institutes of Health Research (PJT-180525) and by Gilead in an investigator iniative study. K.M.C. was funded by NIDDK-sponsored Hepatitis B Research Network (UO-1DK082866 and 3P30DK050306-21S1).

**Resource Tables**

| **REAGENT or RESOURCE** | **SOURCE** | **IDENTIFIER** |
|---|---|---|
| **Antibodies: Flow cytometry** | | |
| Mouse anti-human monoclonal CD68-PE/Cy7 (Y1/82A) | BD Biosciences | Cat# 565595, RRID: AB_2739298 |
| Mouse anti-human monoclonal HLA-DR_BV605 (L243) | BioLegend | Cat#307640, RRID: AB_2561913 |
| Mouse anti-human monoclonal CD86_BV650 (2331) | BD Biosciences | Cat# 563412, RRID: AB_2744456 |

| | | |
|---|---|---|
| Mouse anti-human monoclonal CD40_BV711 (5C3) | BD Biosciences | Cat# 563397, RRID: AB_2738181 |
| Mouse anti-human monoclonal CD206_APC (19.2) | BD Biosciences | Cat# 561763, RRID: AB_398476 |
| Mouse anti-human monoclonal CD209_FITC (DCN46) | BD Biosciences | Cat# 561764, RRID: AB_394122 |
| Mouse anti-human monoclonal CD16_APC/Cy7 (3G8) | BD Biosciences | Cat# 560195, RRID: AB_1645466 |
| **Chemical, peptides, and recombinant proteins** | | |
| Phosphate buffered saline | ThermoFisher Scientific | Cat# 14190-144 |
| RPMI Medium 1640 | ThermoFisher Scientific | Cat# 22400-097 |
| Human Serum | Sigma-Aldrich | Cat# H3667-100mL |

| | | |
|---|---|---|
| Lymphoprep | Stemcell | Cat# 07851 |
| KnockOut Serum | ThermoFisher Scientific | Cat# 10828-028 |
| GlutaMAX (100x) | ThermoFisher Scientific | Cat# 35050-061 |
| Penicillin/ Streptomycin | Lifetech | Cat# 15070063 |
| EDTA | Sigma-Aldrich | Cat# 03690-100ML |
| Bovine serum Albumin | Multicell | Cat# 800-095-CG |
| Saponin | Sigma-Aldrich | Cat# 84510-100G |
| Sodium azide | Sigma-Aldrich | Cat# S2002-100G |
| Paraformaldehyde | Sigma-Aldrich | Cat# 158127-100G |
| Cytofix/ cytoperm | BD Biosciences | Cat# 51-2090KZ |
| Fixable Viability Dye eFluor 450 | eBiosciences | Cat# 65-0863-14 |
| TaqMan fast advanced master mix | Applied Biosystems | Cat# A44360 |

| | | |
|---|---|---|
| Armadillo 384 well PCR plates | Applied Biosystems | Cat# AB2384Y |
| Red Blood Cell Lysis Buffer 10x | BioLegend | Cat# 420301 |
| MCSF | Goldbio | Cat# 1120-09-100 |
| IFN-Beta1b | Goldbio | Cat# 1160-05-2 |
| IFN-Gamma | Goldbio | Cat# 1160-06-20 |
| ApoE3 | Sigma-Aldrich | Cat# SRP4696-500UG |
| IL-10 | Goldbio | Cat# 1110-10-2 |
| IL-4 | Goldbio | Cat# 1110-04-5 |
| **Critical commercial assays** | | |
| CD14+ microbeads | Miltenyi | Cat# 130-050-201 |
| RNeasy mini kit | Qiagen | Cat# 74106 |
| High-capacity cDNA Reverse transcription kit | Applied Biosystems | Cat# 43-688-14 |
| Legendplex™ human inflammation | BioLegend | Cat# 740809 |

| | | |
|---|---|---|
| panel 1 (13-plex) with V-bottom plate | | |
| Custom Luminex Bead Assay | R&D Systems | N/A |
| scRNAseq 5' v2 | 10x genomics | N/A |
| **Oligonucleotides: TaqMan qPCR probe information** | | |
| C1QA | ThermoFisher Scientific | Hs00381122_m1 |
| VCAN | ThermoFisher Scientific | Hs00171642_m1 |
| FCGR3A | ThermoFisher Scientific | Hs02388314_m1 |
| IFI27 | ThermoFisher Scientific | Hs01086370_m1 |
| IFIT3 | ThermoFisher Scientific | Hs00155468_m1 |
| IL18 | ThermoFisher Scientific | Hs01038788_m1 |
| NR1H3 | ThermoFisher Scientific | Hs00172885_m1 |

| | | |
|---|---|---|
| ZFP36L1 | ThermoFisher Scientific | Hs00245183_m1 |
| MAFB | ThermoFisher Scientific | Hs00534343_s1 |
| CETP | ThermoFisher Scientific | Hs00163942_m1 |
| **Software and algorithms** | | |
| Cell Ranger | 10x Genomics | https://support.10xgenomics.com/singlecell-gene-expression/software/pipelines/latest/what-is-cell-ranger |
| R version 4.0.3 | The R project | https://www.r-project.org/ |
| Seurat | (Stuart et al., 2019) | https://satijalab.org/seurat/ |
| Scran | (Lun et al., 2016) | https://bioconductor.org/packages/release/bioc/html/scran.html |
| pheatmap | N/A | https://rdrr.io/cran/pheatmap/ |
| ggplot2 | (Wickham, 2016) | https://ggplot2.tidyverse.org |
| Fgsea | (Korotkevich et al., 2021) | https://bioconductor.org/packages/release/bioc/html/fgsea.html |

| | | |
|---|---|---|
| Monocle 2 | (Qiu, et al., 2017) | http://cole-trapnell-lab.github.io/monocle-release/docs/ |
| Nichenet | (Browaeys et al., 2019) | https://github.com/saeyslab/nichenetr |
| GraphPad Prism 9 | GraphPad | https://www.graphpad.com/ |
| FlowJo | FlowJo, LLC | https://www.flowjo.com |
| xPONENT | Luminex Corporation | https://www.luminexcorp.com/xponent/#overview |
| **Additional Databases** | | |
| Healthy Liver scRNAseq data 3' V2 | (MacParland et al., 2018) | GEO accession GSE115469 |
| Cirrhotic Liver scRNAseq data 3' V2 | (Ramachandran et al., 2019) | GEO accession GSE136103 |
| Long term NUC scRNAseq data | | PMID: 36728214 |

**Supplemental information**

**Suppl. table 1. Clinical characteristics of patients with chronic hepatitis B that were included in this study.**

| | Mean | Range |
|---|---|---|
| Age [years] | 45.2 | 29–64 |
| Male/Female [% of all patients] | 67/33 | |
| ALT at Baseline [* ULN] | 8.9 | 1.1–21.8 |
| HBV DNA at screening [IU/mL] | $3.09 \times 10^7$ | $2.73 \times 10^4$–$9.97 \times 10^7$ |
| HBeAg (+) at Baseline [% of all patients] | 44% | |

**Suppl. Figure 1. Cell populations extracted from FNAs and sequenced using scRNAseq were consistent and equally distributed across patients. A)** UMAP dimensionality reduction corresponding to the one in Figure 1D for each individual patient across 3 timepoints (baseline, week12 and week24). UMAP dimensionality reduction identified 30 clusters. **B)** Each cluster was annotated and assigned to specific cell types using differential gene expression. All selected genes have an adjusted p value < 0.05. FNAs, fine needle aspirates; scRNAseq, single cell RNA sequencing.

**Suppl. Figure 2. Characterization and annotation of liver macrophages and CD8 T cells by IMC. A)** IMC individual marker assessment for characterization of liver macrophages (top row) and CD8 T cells (bottom row) from the inflamed liver of a CHB patient. Portal regions are outlined in yellow dotted lines. **B)** Voltage determination of positive and negative populations for annotation of liver macrophages and CD8 T cells using individual markers.

**Suppl. Figure 3. IL-10 stimulation is needed for CD16 upregulation while not impairing the inflammatory potential of the *in vitro* differentiated macrophages. A)** Real-time qPCR analyses of relative fold change for mRNA expression on differentiated Macs using individual and combined ligands predicted by NicheNet. **B)** NicheNet analyses predicted IL-10 has the highest regulatory potential score for upregulation of CD16 (*FCGR3A*) at baseline, compared to week 24 in the iMacs. **C)** The interferon stimulated-gene response and IL-18 is not dampened by IL-10 while preserving CD16 (FCGR3A) expression. “Combined” ligands include MCSF, IFN-β, IFN-γ and Apolipoprotein E (ApoE). *P*values determined by repeated-measures one-way ANOVA, (*$P$ <0.05, **$P$ <0.005, ***$P$ <0.001). Results showed are representative of **n=**5 experiments. MCSF, macrophage colony-stimulating factor; IFN, interferon; iMacs, inflammatory macrophages; Macs, macrophages.

**Suppl. Figure 4. The *in vitro* differentiated macrophages resemble canonical M1 macrophages and have the potential to drive inflammation. A)** Protein expression of canonical M1 (pro-inflammatory) and M2 (anti-inflammatory) Macs in different types of *in vitro* differentiated Macs. **B)** Inflammatory cytokines released by different types of *in vitro* differentiated Macs, **n=** 6. MFI, median fluorescence intensity; M0, MCSF only stimulated macrophages; MCSF, macrophage colony-stimulating factor; iMacs, inflammatory macrophages; Macs, macrophages; TLR, toll-like receptor. “Induced iMac” ligands include M-CSF, IFN-β, IFN-γ, Apolipoprotein E (ApoE) and IL-10. *P*values determined by repeated-measures one-way ANOVA, (**P* <0.05, ***P* <0.005, ****P* <0.001). Results showed are representative of **n=**5 experiments, unless otherwise specified.

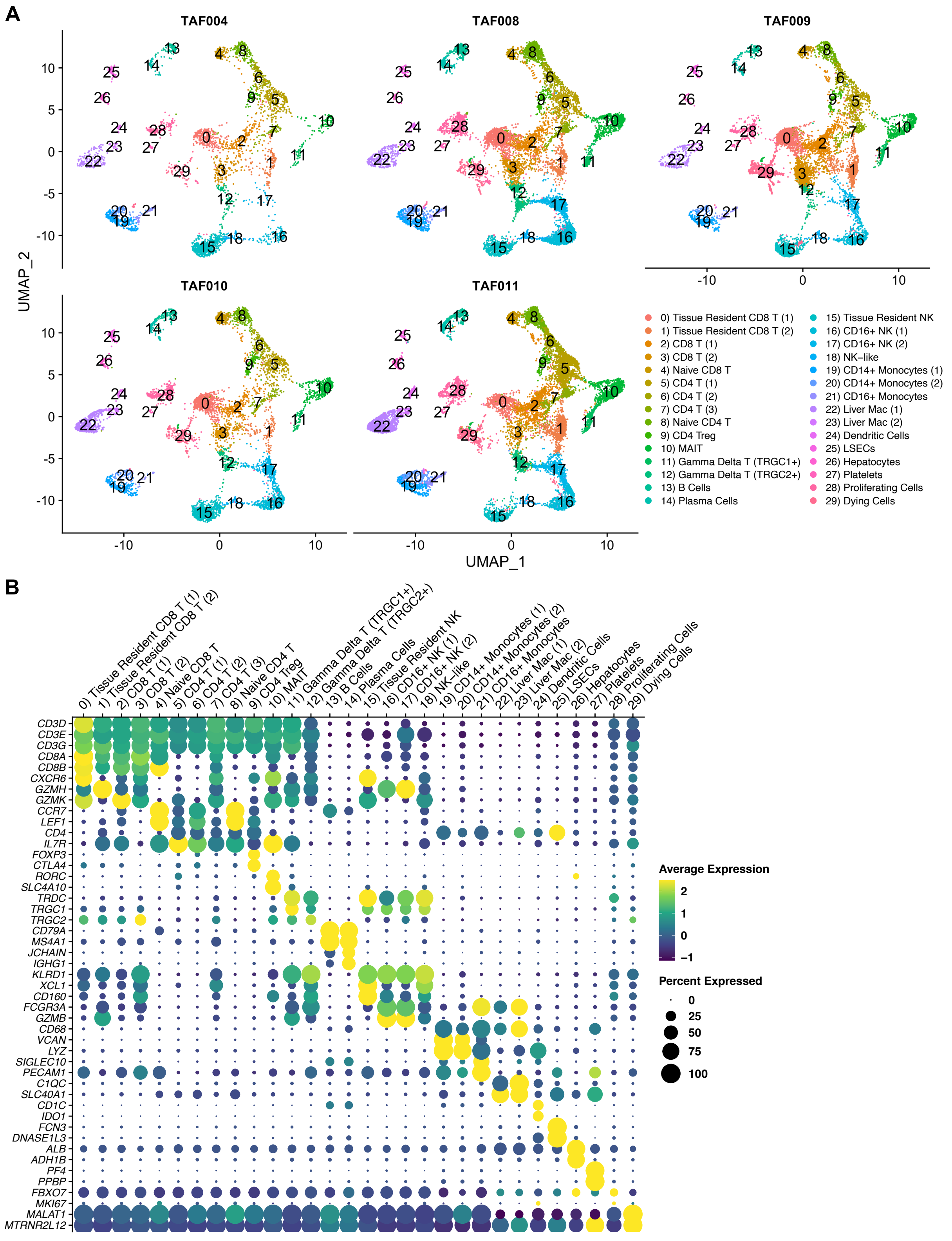


**Suppl. Figure 1. Cell populations extracted from FNAs and sequenced using scRNAseq were consistent and equally distributed across patients. A)** UMAP dimensionality reduction corresponding to the one in Figure 1D for each individual patient across 3 timepoints (baseline, week12 and week24). UMAP dimensionality reduction identified 30 clusters. **B)** Each cluster was annotated and assigned to specific cell types using differential gene expression. All selected genes have an adjusted p value < 0.05. FNAs, fine needle aspirates; scRNAseq, single cell RNA sequencing.

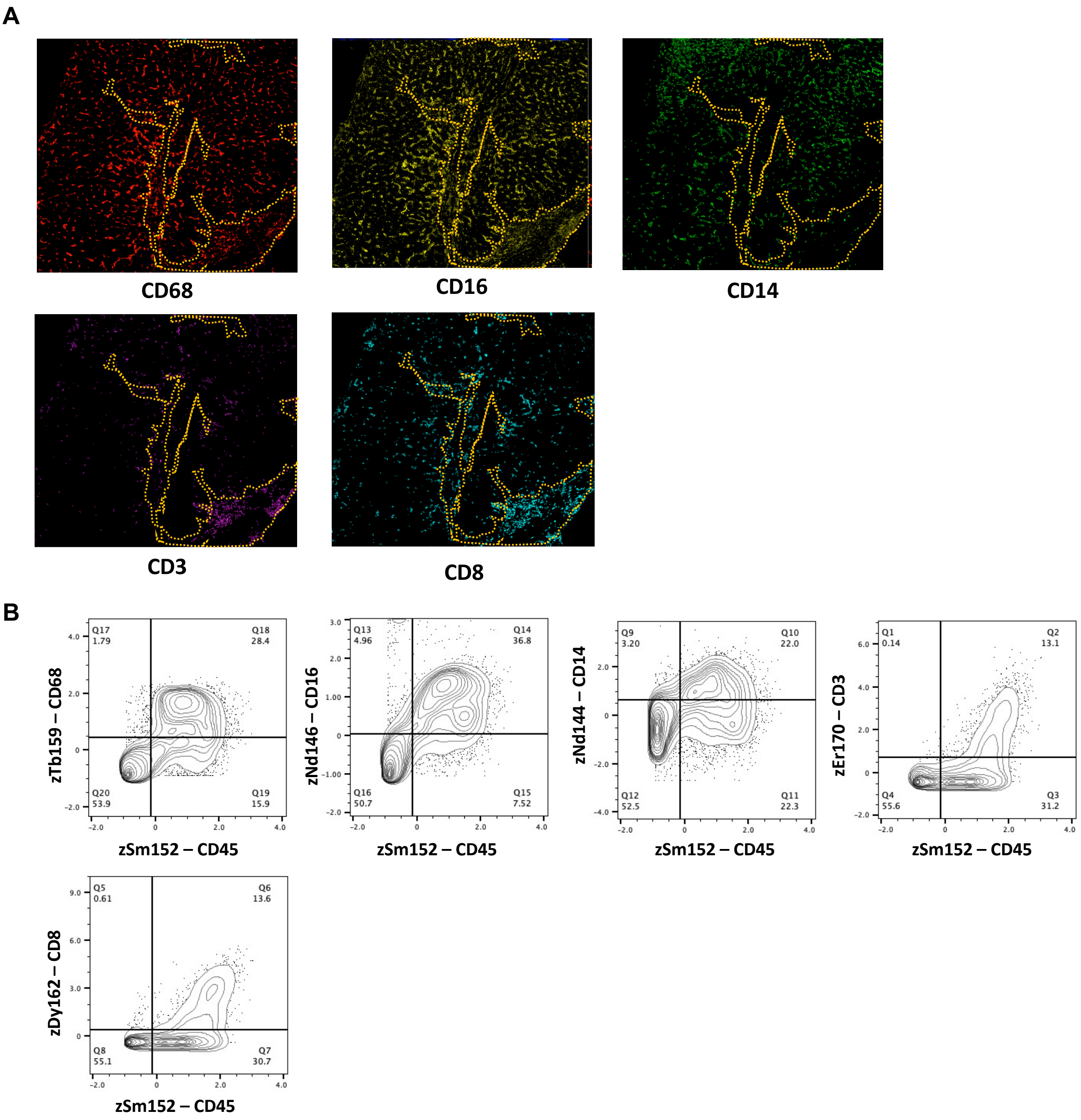


**Suppl. Figure 2. Characterization and annotation of liver macrophages and CD8 T cells by IMC. A)** IMC individual marker assessment for characterization of liver macrophages (top row) and CD8 T cells (bottom row) from the inflamed liver of a CHB patient. Portal region outlined in yellow dotted lines. **B)** Voltage determination of positive and negative populations for annotation of liver macrophages and CD8 T cells using individual markers.

**A**

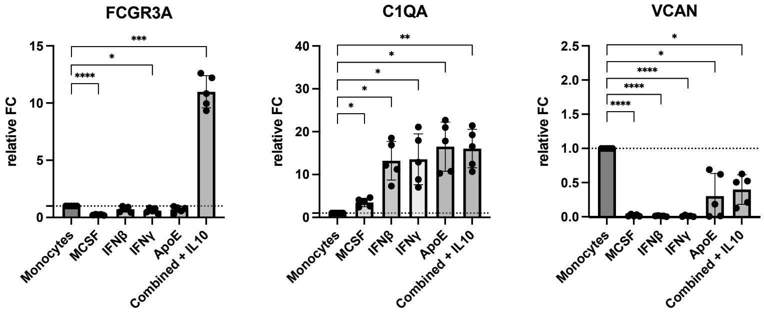


**B**

| Ligand | Regulatory potential score for CD16 in iMacs |
|---|---|
| IL10 | 0.004162998 |

**C**

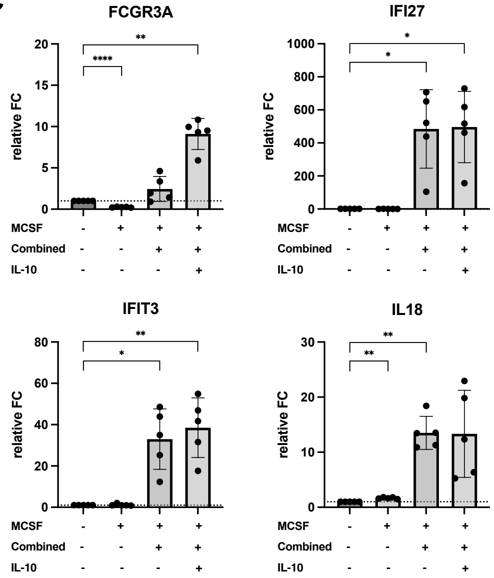


**Suppl. Figure 3. IL10 stimulation is needed for CD16 upregulation while not impairing the inflammatory potential of the *in vitro* differentiated macrophages. A)** Real-time qPCR analyses of relative fold change for mRNA expression on differentiated macrophages using individual and combined ligands predicted by NicheNet. **B)** NicheNet analyses predicted IL10 has the highest regulatory potential score for upregulation of CD16 (*FCGR3A*) at baseline, compared to week 24 in the iMacs. **C)** The interferon stimulated-gene response and IL18 is not dampened by IL10 while preserving CD16 (FCGR3A) expression. “Combined” ligands include MCSF, IFN-β, IFN-γ and Apolipoprotein E (ApoE). *P*values determined by repeated-measures one-way ANOVA, (**P* <0.05, ***P* <0.005, ****P* <0.001 ). Results showed are representative of **n**=5 experiments. MCSF, macrophage colony-stimulating factor; IFN, interferon; iMacs, inflammatory macrophages.

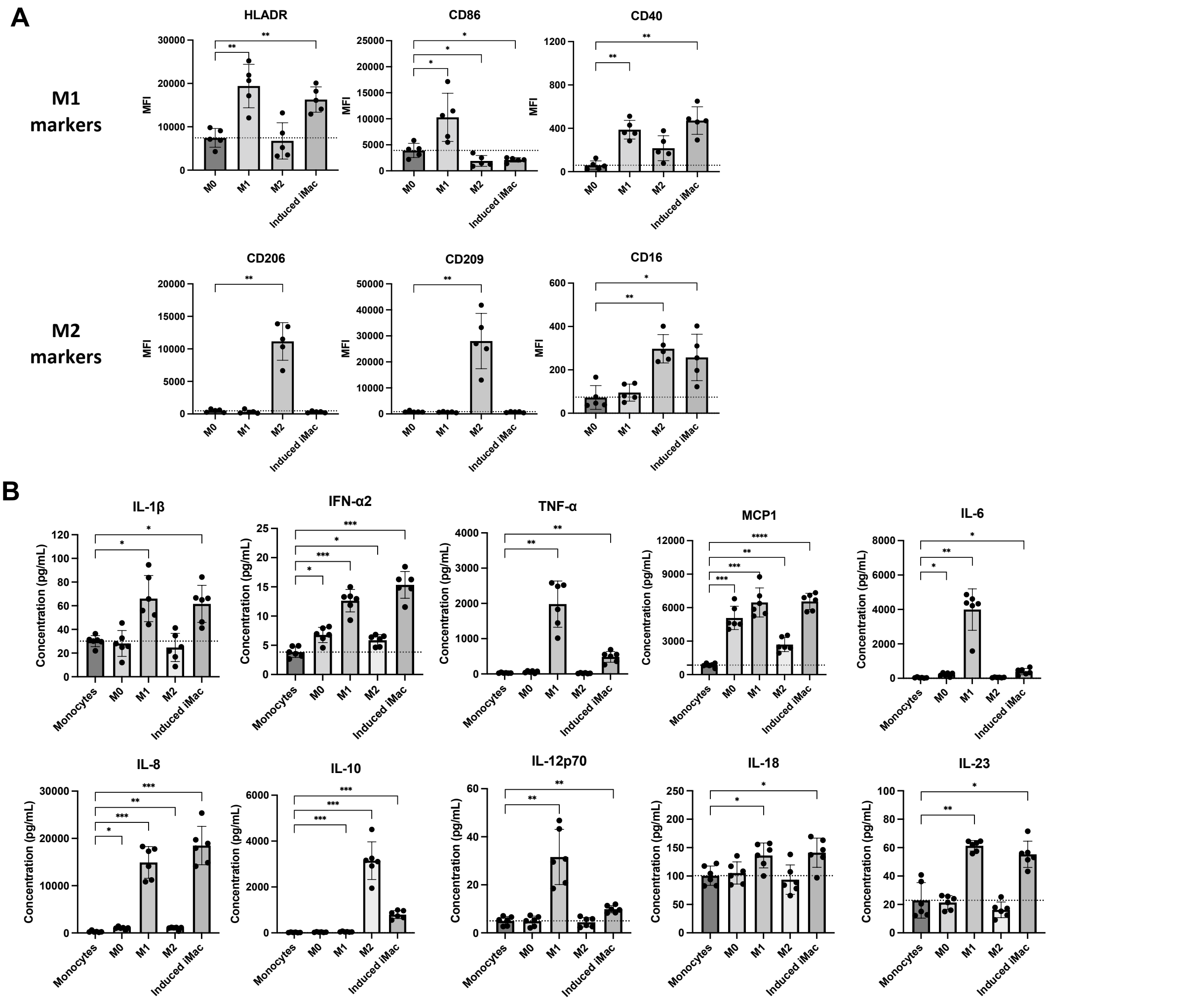


**Suppl. Figure 4. The *in vitro* differentiated macrophages resemble canonical M1 macrophages and have the potential to drive inflammation. A)** Protein expression of canonical M1 (pro-inflammatory) and M2 (anti-inflammatory) macrophages in different types of *in vitro* differentiated macrophages. **B)** Inflammatory cytokines released by different types of *in vitro* differentiated macrophages, **n=** 6. MFI, median fluorescence intensity; M0, MCSF only stimulated macrophages; MCSF, macrophage colony-stimulating factor; iMacs, inflammatory macrophages; Macs, macrophages; TLR, toll-like receptor. "Induced iMac" ligands include M-CSF, IFN-β, IFN-γ, Apolipoprotein E (ApoE) and IL-10. *P*values determined by repeated-measures one-way ANOVA, (**P* <0.05, ***P* <0.005, ****P* <0.001 ). Results showed are representative of **n=**5 experiments, unless otherwise specified.
